# Distinct and cooperative roles of host and tumor Osteopontin in colorectal cancer liver metastasis

**DOI:** 10.64898/2026.02.19.706899

**Authors:** Patrick Czabala, Yang Zhao, John D. Klement, Priscilla S. Redd, Dakota Poschel, Kristen Carver, Kendra Fick, Zainab Tiamiyu, Martina Zoccheddu, Patricia V. Schoenlein, Jennifer Waller, Huidong Shi, Kebin Liu

## Abstract

Osteopontin (OPN) is a secreted phosphoprotein implicated in colorectal cancer liver metastasis (CRCLM), yet the distinct spatial contributions of host-and tumor-derived OPN in driving this disease remain unclear. Using a 2 x 2 genetic knockout mouse model targeting OPN in host and tumor compartments, combined with spatial transcriptomics, we investigated compartment-specific OPN functions in CRCLM. Tumor-derived OPN promotes tumor proliferation through MEK/ERK signaling. Host OPN licenses monocyte-to-macrophage differentiation, while tumor OPN polarizes macrophages towards an M2-like state. Both host and tumor OPN suppress T cells in the tumor microenvironment, whereas loss of host OPN reveals an interferon-driven, anti-tumor niche. Translational studies using OPN-blockade immunotherapy in syngeneic and patient-derived xenograft mouse models reduced tumor burden and enhanced T cell infiltration. Together, these findings redefine the OPN-myeloid paradigm in CRC and nominate OPN as a potential therapeutic target.

## Introduction

Colorectal cancer (CRC) is the second most common cause of cancer death in the United States^1^. Although a minority of CRCs (10-15%) exhibit microsatellite instability-high (MSI-H) status and respond well to PD-(L)1 blockade, most disease (85-90%) is microsatellite stable and mismatch repair-proficient (MSS), a subtype that responds poorly to immune checkpoint inhibition (ICI)^2,3^. Compounding the clinical challenge of MSS CRC management, up to 50% of patients with CRC will develop liver metastasis (CRCLM), which is difficult to resect and accounts for approximately 70% of metastatic spread in CRC due to hematogenous dissemination through the portal system^4,5,6^. Liver metastasis is an independent predictor of poor prognosis in CRC and is associated with substantially reduced responsiveness to PD-(L)1 blockade compared to metastasis at other sites^7–9^.

Despite these poor clinical outcomes, both MSS and MSI-H CRC frequently contain abundant cytotoxic T lymphocytes (CTLs), suggesting that immune exclusion alone does not fully explain therapeutic resistance^10–12^. Notably, PD-L1 expression is often weak or absent on tumor epithelial cells in MSS CRC, indicating that failure of ICI therapy may be due to PD-L1-independent mechanisms^13,14^. Emerging evidence suggests that the liver metastatic environment imposes unique constraints on antitumor immunity^15^. Therefore, there is an urgent unmet need to understand the mechanisms that promote liver metastasis and to identify new avenues for treatment, particularly in MSS disease.

Among biomarkers of CRC progression, Osteopontin has become a major target of recent investigation^16–18^. Osteopontin (OPN), encoded by *SPP1*, is a secreted phosphoprotein that is expressed by tumor cells and various host and immune cells. OPN binds to CD44 and multiple integrin heterodimers, modulating extracellular matrix interactions, cell adhesion, and intracellular signaling^19^. We and others have shown that OPN suppresses T cell function via CD44 and promotes macrophage recruitment through CD44 and integrin signaling^20–23^. In multiple cancers, including CRC, OPN promotes progression and metastasis^24–33^. Notably, CRCLM tumors are enriched for *SPP1*-high, M2-like tumor-associated macrophages (TAMs), a population recurrently associated with poor clinical outcomes^34–36^. This population of SPP1-hi macrophages has also been shown to mediate ICI resistance in castration-resistant prostate cancer^37^. OPN is also expressed by tumor cells and has recently been shown to maintain mesenchymal cell fate in pancreatic cancer, underscoring a broad role in regulating tumor cell fate^38,39^. OPN has been proposed as a therapeutic target; however, systemic approaches that globally neutralize OPN are likely to affect both host-and tumor-derived pools^40,41,42^. The relative contributions of tumor-derived versus host-derived OPN in CRCLM have not been directly investigated to this point.

Recent advancements in spatial transcriptomics enable single-cell resolution and gene expression mapping within tumors. Here, we leveraged CosMx spatial transcriptomics and genetic knockout mouse models to interrogate the compartment-specific functions of OPN in liver metastasis, revealing distinct tumor-and host-derived roles in shaping the tumor microenvironment in CRCLM. Targeting OPN may unlock an anti-tumor immune niche associated with responsiveness to PD-(L)1 blockade.

## Results

### Tumor-and host-derived OPN promote colon cancer liver metastasis

Analysis of colorectal cancer (CRC) datasets from The Cancer Genome Atlas (TCGA; COAD/READ) revealed that high OPN expression is associated with reduced overall survival time in patients with colon cancer (Fig. 1A). Consistent with this observation, OPN expression level increases with disease progression and was highest in patients with stage IV CRC (Fig. 1B). To define cellular sources of OPN in CRC, we next analyzed publicly available single-cell RNA sequencing (scRNAseq) datasets from human colon cancer patients. OPN expression was highest in macrophages in liver metastases (CRCLM) (Fig. 1C). Notably, OPN was also detected in tumor cells, with significantly higher expression in liver metastatic tumor cells compared to primary tumor cells (Fig. 1D).

**Figure 1.**
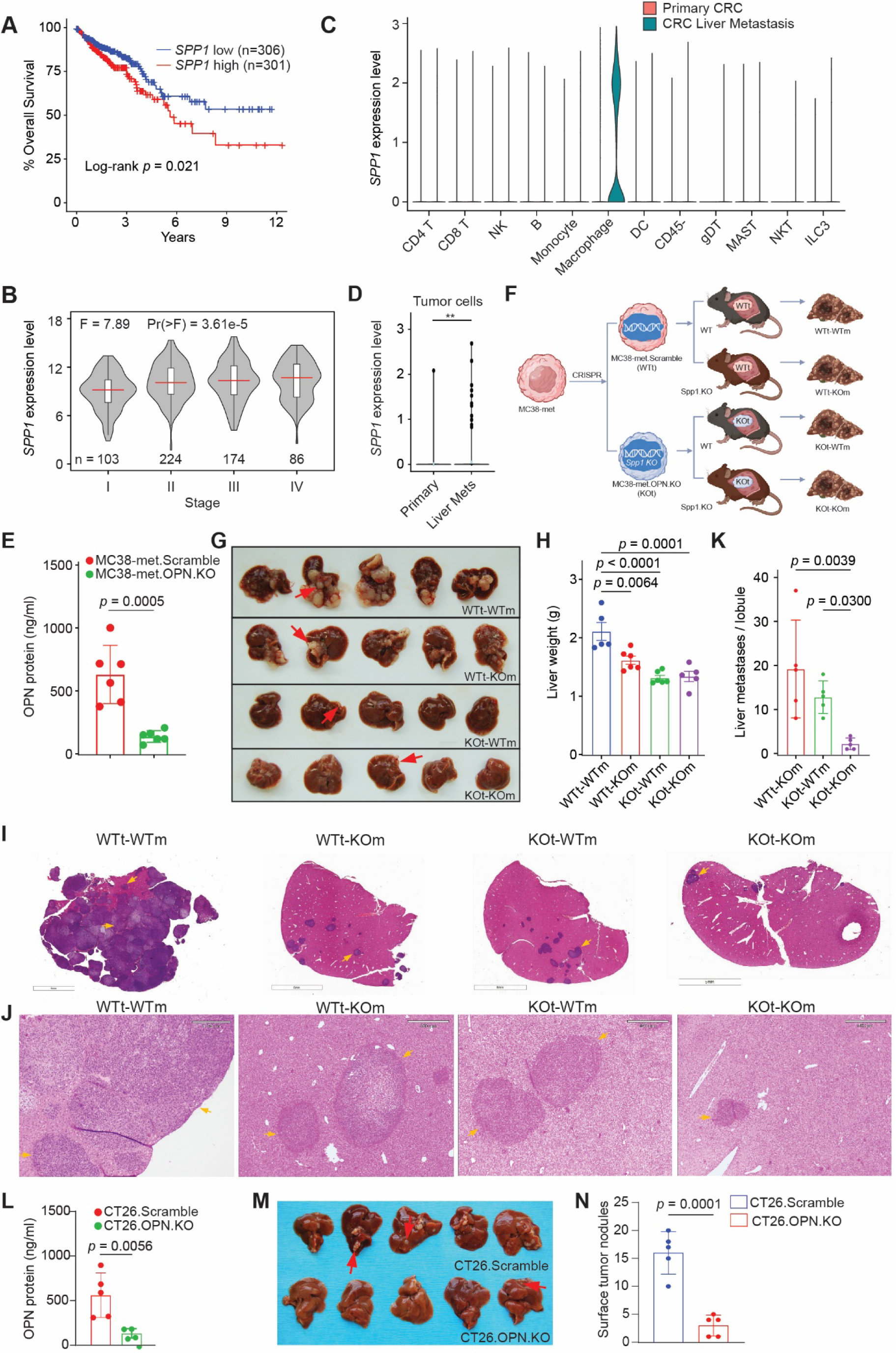
Tumor-and host-derived Osteopontin drive colon cancer liver metastasis. (A) Kaplan-Meier analysis (TCGA COAD/READ) demonstrates that high *SPP1* expression in primary tumors correlates with shorter overall survival (50% cutoff; log-rank test). (B) Violin plots reveal increased *SPP1* expression with advancing tumor stage (TCGA COAD/READ; one-way ANOVA). (C) scRNAseq from colon cancer patients. *SPP1* expression is highest in the macrophages of liver metastasis. (D) scRNAseq from colon cancer patients. *SPP1* expression is expressed in tumor cells and is higher in liver metastasis. (E) OPN knockout validation in MC38met cells using ELISA. (F) Schematic of 2×2 liver metastasis model: MC38-met WT or OPN-KO cells administered into WT or OPN-KO mice (n = 5-6/group). (G) Representative images of liver metastases. (H) Liver weights corresponding to (F & G). One-way ANOVA with Dunnett’s multiple comparisons using WTt-WTm as the control group. (I) Representative H&E-stained liver sections from (G) showing metastatic nodules (arrows). (J) Magnified views of (I). (K) Quantification of microscopic metastases per lobule. One-way ANOVA with Holm-Sidak multiple comparisons of the single KO groups to the KOt-KOm group. (L) OPN knockout validation in CT26 cells using ELISA. (M) Liver metastases in BALB/c mice administered CT26 WT or OPN-KO cells. Metastatic nodules indicated by yellow arrows. (N) Quantification of surface liver metastases from (M). Data presented as mean ± s.d., each dot = one biological replicate. Two-tailed Student’s t tests were used for all other comparisons.

These findings indicate that OPN is preferentially enriched in tumor cells and tumor-associated macrophages (TAMs) in CRCLM, suggesting that OPN may promote liver metastatic disease. To test this hypothesis, we knocked out *Spp1* in the murine colon tumor cell line, MC38-met (Fig. 1E). WT (WTt) and OPN-deficient (KOt) tumor cells were transplanted to C57BL/6J mice (WTm) and B6.129S6(Cg)-*Spp1^tm1Blh^*/J (hereafter referred to as OPN-KO; KOm) mice using an experimental liver metastasis model involving intrasplenic injection followed by splenectomy^43^, mimicking the portal venous route of metastasis (Fig. 1F). Loss of OPN in tumor cells, host tissue, or both resulted in a significant reduction in liver metastases burden as measured by gross liver weight (Fig. 1G & H). Histological analysis confirmed significantly fewer microscopic tumor nodules formed when OPN KO tumor cells were implanted into *Spp1* KO mice (KOt-KOm) compared with all other groups (Fig. 1I-K).

To validate these observations in an independent model, *Spp1* was then knocked out in a second murine colon tumor cell line, CT26 (Fig. 1L). WT or OPN-KO CT26 tumor cells were surgically transplanted into syngeneic BALB/c mice to establish liver metastasis. As observed in the MC38-met model, knocking out OPN significantly liver metastatic burden (Fig. 1M & N).

Together, these genetic loss-of-function studies demonstrate that both tumor-and host-derived OPN contribute to CRCLM. However, these experiments do not resolve how OPN from distinct compartments shapes tumor cell states, immune phenotypes, or cell-cell interactions within the metastatic microenvironment. Given that OPN is a secreted ligand whose signaling depends on spatial proximity between cells that produce or respond to it, we turned to spatial transcriptomics to resolve its source-specific functions.

### Spatial transcriptomics reveals source-specific OPN alters cell phenotypes and local microenvironments in colon tumor liver metastasis

We employed spatial transcriptomics to characterize how OPN spatially modulates cellular phenotypes and interactions within CRCLM. Using liver metastasis samples from the 2×2 OPN knockout experiment (Fig. 1F), we applied the CosMx Mouse Universal Characterization panel on the Spatial Molecular Imager (SMI) platform (Fig. 2A). The panel simultaneously detected 1,000 selected mRNA transcripts, three cell-surface proteins, and a nuclear stain at subcellular resolution on FFPE slides. A total of 544 fields of view (FOVs) in the tumor nodule areas were selected in the four experimental groups. Extensive single-cell segmentation was performed (Fig. S1A), yielding a total of 1,168,484 single cells.

**Figure 2.**
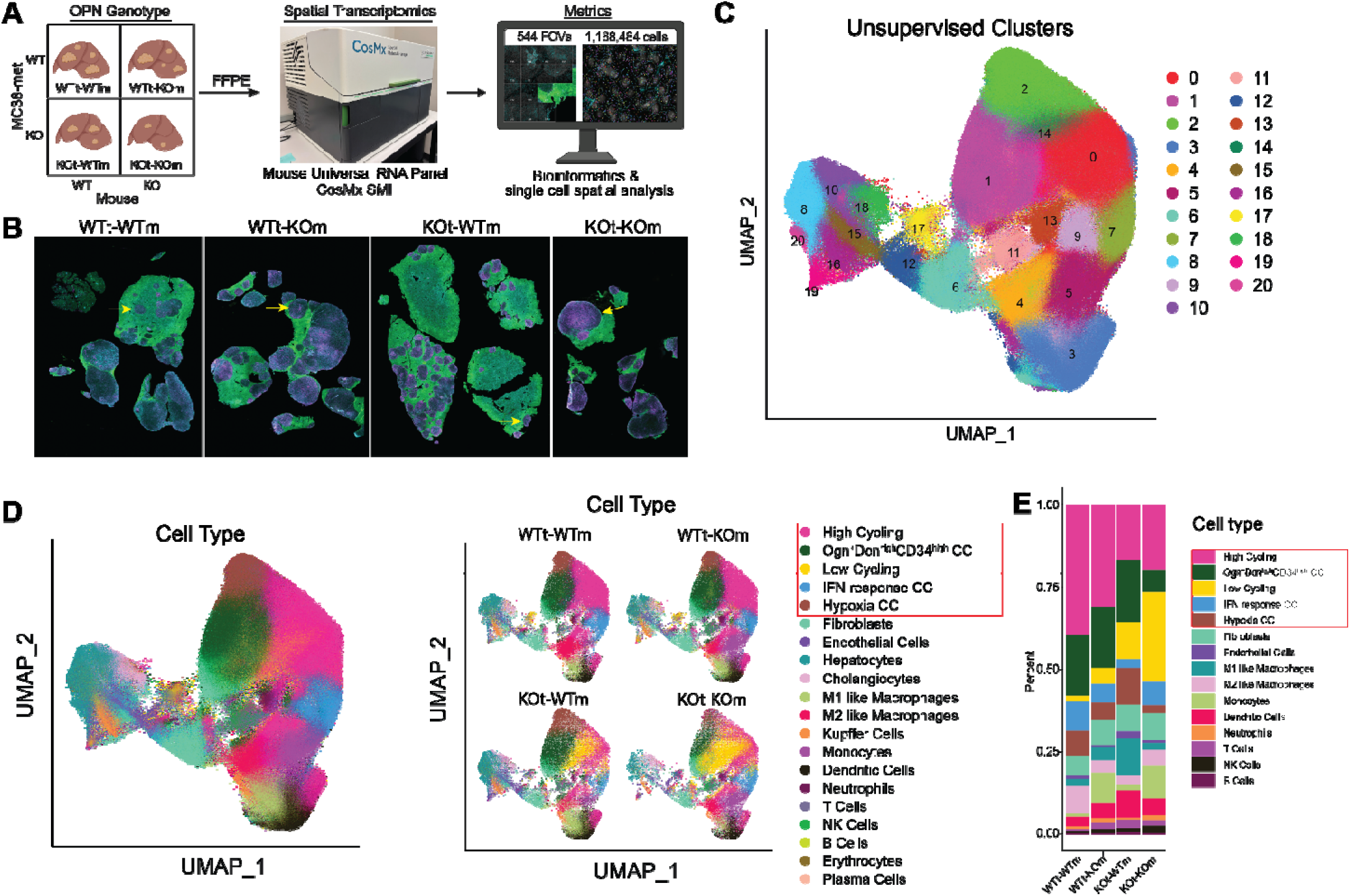
Tumor-and host-derived OPN regulate cell phenotypes in the tumor microenvironment. (A) Schematic of CosMx spatial transcriptomics experiment (B) Immunofluorescence composite image of the samples from the CosMx experiment in (A). Pan-CK (green), CD298/B2M (blue), CD45 (pink), DNA (grey) (C) Unsupervised clustering of the integrated spatial transcriptomics data identified 21 cell populations, visualized by UMAP. (D) Overall UMAP of high-resolution annotation of cells (left) and split by sample (right). Cell types are indicated in the legend on the right. A box has been added to the legend to emphasize the tumor cell subtypes. (E) Bar plot of proportion of cells inside the tumor nodules only.

Unsupervised clustering identified 21 cell populations (Fig. 2C, 2D, & S1B). Cell-typing annotated the non-cancer cells into 15 subpopulations, including 8 immune cell clusters and 7 stromal and parenchymal cell clusters (Fig. 2D & E). Published colon tumor cell subpopulation signatures were used to perform reference mapping to annotate the five tumor cell subpopulations: high-cycling, low-cycling, Ogn^+^dcn^hi^CD34^hi^ (stem-like), IFN-response, and hypoxia cancer cells (Fig. 2D)^44^. To quantify the frequency of cell types in the tumors, we created bar plots of cells located inside nodules only, excluding normal liver parenchyma that surrounded the tumors (Fig. 2E & 3A).

The spatial location provided by spatial transcriptomics enables neighborhood/niche analysis that groups localized microenvironments based on groups of cells that are co-localized. We performed niche analysis and identified eight tumor niches (T1-T8) and two liver niches (L1 and L2) (Fig. 3C). We plotted representative FOVs that were comprised of essentially a single niche (Fig. 3F, S2C) and observed that distinct cell types defined each niche (Fig. 3E). Niche T1 and T2 were dominated by high-cycling tumor cells and low infiltration of immune cells. Niche T3 exhibited a high proportion of hypoxia tumor cells, while T4 contained many stem-like cancer cells. Low-cycling cells predominate in niche T5. Niches T6 and T7 represent the two immune-response niches: in T6, there are high monocytes, dendritic cells (DCs), and T cells, while T7 was characterized by M1-like macrophages, DCs, and the highest proportion of T cells in any niche. Finally, niche T8 represented the pseudocapsules surrounding the tumor nodules (Fig. 3B). Notably, we observed heterogeneous enrichment of niches across different samples (Fig. 3C & D). High-cycling niches T1 and T2 were highest in the WTt-WTm sample, while in the globally OPN-deficient KOt-KOm, the low-cycling niche T5 was most abundant, hinting at the role of OPN in driving tumor cell proliferation. The immune active niches T6 and T7 were essentially absent in WTt-WTm tissue. T6 was enriched in OPN-deficient hosts (WTt-KOm and KOt-KOm), while loss of tumor OPN led to an increase in T7 (KOt-WTm) (Fig. 3C & D).

**Figure 3.**
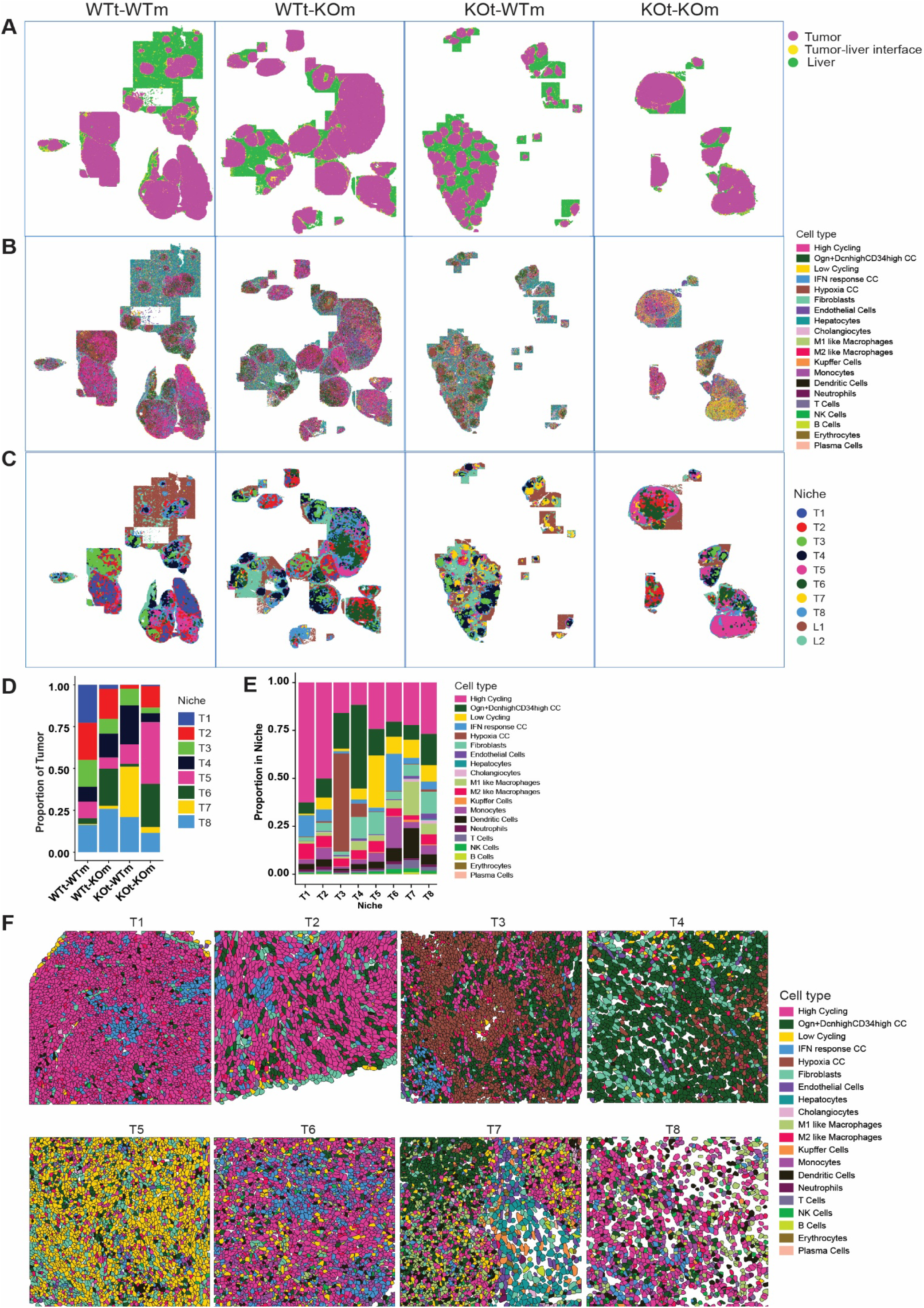
Eight distinct tumor niches exhibit distinct cell-type proportions and are differentially regulated by tumor-and host-derived OPN. (A-C) Spatial maps showing liver and tumor regions (A), cell-type distributions (B), and tumor and liver niches (C). (D-E) Bar plots of proportions of cell types (D) and niches (E) within tumor nodules across the four experimental groups. (F) Representative FOVs enriched for a single niche, colored by cell type.

Spatial transcriptomic profiling revealed widespread effects of tumor-and host-derived OPN across the tumor and immune compartments. We focused subsequent analyses on four mechanistic programs that are biologically and therapeutically relevant in CRCLM. Thse include: (i) Autocrine OPN signaling in tumor cells drives proliferation, (ii) host-and tumor-derived OPN differentially regulate monocyte differentiation and polarization, (iii) OPN suppresses T cells in the TME, and (iv) IFN-response programs dominate in the tumors of OPN-deficient hosts.

### Autocrine tumor-derived OPN drives tumor proliferation through MAPK signaling

Based on the observation that the high cycling tumor niches T1 and T2 were most abundant in the WTt-WTm group, whereas the low-cycling niche T5 was enriched in the KOt-KOm group, we investigated how OPN influences tumor proliferation. Over 50% of the cells in T1 are high-cycling tumor cells (Fig. 4A). To identify the signals driving proliferation in these cells, we applied CellChat analysis to a representative T1 FOV and found that the strongest receptor-ligand interactions were autocrine interactions among high cycling tumor cells (Fig. 4B & C, S3A-D). Examining specific networks revealed that the SPP1 (OPN) network was a significant contributor to these autocrine interactions (Fig. 4E & F, & S3C-D).

**Figure 4.**
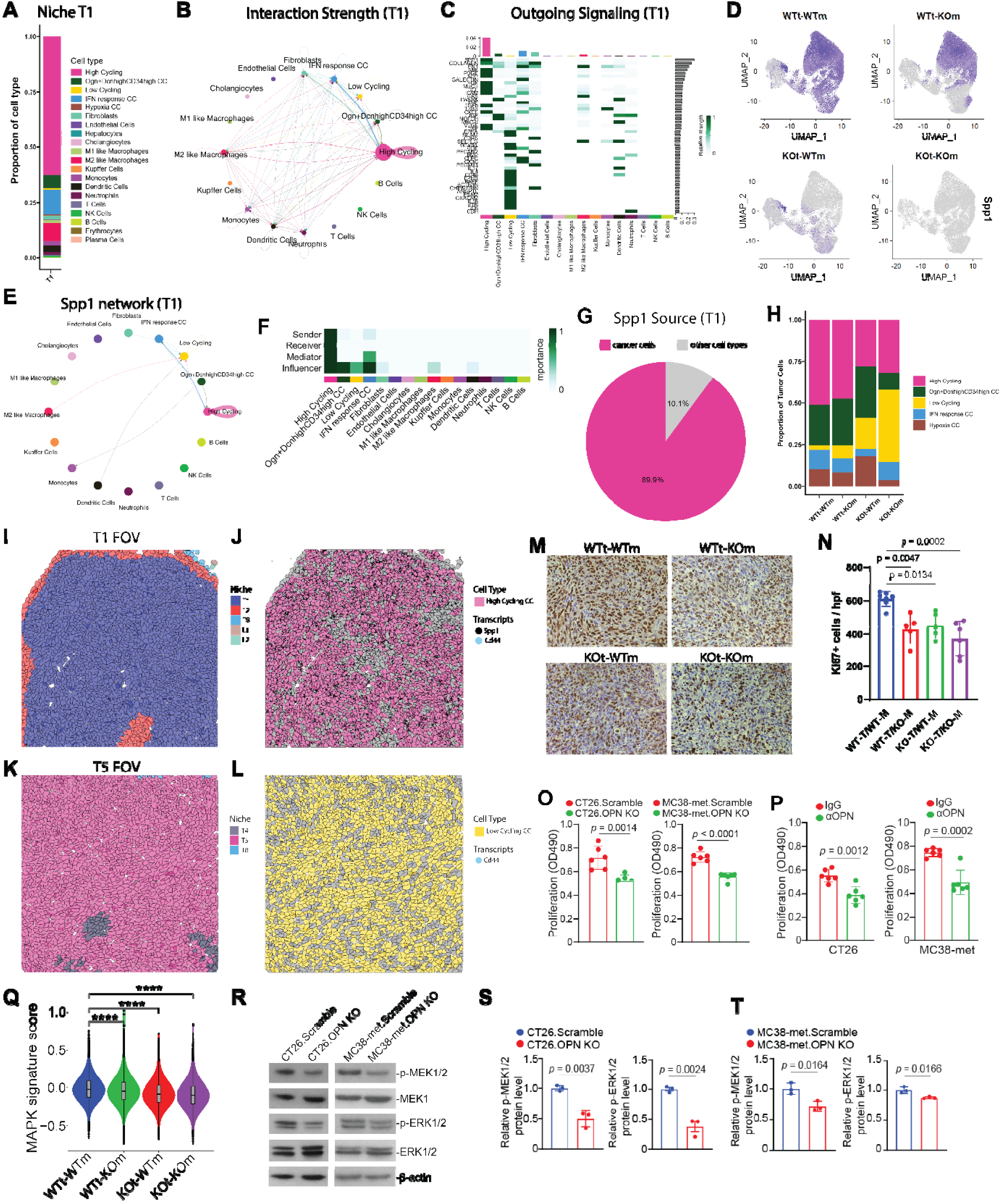
Tumor-derived OPN drives autocrine tumor cell proliferation via the MEK/ERK pathway A) Bar plot of cell-type composition in niche T1, highlighting high-cycling cancer cells as the predominant subset (B) Circular plot of cell-cell communication analysis of representative T1 Field of View (FOV), using CellChat based on known ligand–receptor pairs; Nodes represent cell types. Node size reflects the total communication strength of the cell population. Directed edges represent predicted signaling interactions, with arrows indicating the direction from ligand-expressing sender cells to receptor-expressing receiver cells. (C) Heatmap shows the outgoing signaling patterns of the representative T1 FOV. A gradient of white to dark green indicates low to high expression weight value in the heatmap. (D) UMAP Plots of normalized *Spp1* expression levels across the four experimental groups, indicating primary expression in tumor and myeloid populations. (E) Circular plot of SPP1 receptor-ligand interaction network in representative T1 FOV; connection thickness represents the net aggregate of the SPP1 network. (F) Heatmap of the SPP1 Signaling network displaying relative importance of each cell group ranked according to the computed four network centrality measures. The mediator score is quantified using betweenness centrality, whereas the influencer score is quantified by information centrality. (G) Pie chart of *Spp1* expression in the same FOV, showing tumor cells as the dominant source. (H) Bar plot of cancer cell subtype proportions within tumor nodules. (I-J) Representative T1 FOV colored by niche (I) or high cycling cancer cells (J) with *Spp1* (black) and *Cd44* transcripts (blue). (K-L) Representative FOV of the T5 low-cycling Niche from KOt-KOm colored by niche (K) and low-cycling cancer cells (L) with *Cd44* transcripts (blue). (*Spp1* transcripts are present due to CRISPR translation-blocking KO; removed in the image for clarity) (M-N) Ki-67 IHC in tumor nodules: representative images (I) and quantification of Ki-67+ cells per high-powered field (J; n = 5-7), showing reduced proliferation in OPN-knockout groups relative to WT. (O-P) In vitro proliferation of CT26 and MC38met cells: OPN-KO lines proliferate slower than scramble controls (K, n = 5 biological replicates), and treatment with anti-OPN mAb reduces proliferation relative to IgG control (L, n = 6 biological replicates). Q) Violin plot of MAPK signature gene score in colon cancer cells across the four experimental groups; knockout groups display lower scores than WT. (R-T) Western blot analysis of MEK and ERK phosphorylation: representative blots (R), and quantification of p-MEK1/2 (S) and p-ERK (T).

Feature plots of *Spp1* expression across the four samples showed that OPN is expressed by tumor cells, myeloid cells, and liver parenchymal cholangiocytes (Fig. 4D). Image plots of a representative T1 FOV, composed primarily of niche T1 (Fig. 4I), visually confirmed expression of *Spp1* and *Cd44*, a receptor for OPN, in this niche (Fig. 4J, & S4A-D). Although OPN is expressed by tumor cells, it is even higher in myeloid cells, particularly macrophages in CRCLM patients (Fig. 1C). Quantification of *Spp1* transcripts revealed that, in niche T1, nearly 90% of *Spp1* mRNA originates from tumor cells (Fig. 4G), suggesting that autocrine OPN signaling predominates over myeloid-derived OPN in driving tumor proliferation in this model. Consistent with this, the proportion of low cycling tumor cells was higher in the KOt-WTm group than the WTt-KOm group (Fig. 4H). A representative low-cycling T5 FOV from the KOt-KOm sample illustrates global OPN ablation (Fig. 4K & L). Furthermore, the KOt-KOm group displayed an increase in low-cycling tumor cells relative to the KOt-WTm group (Fig. 4H), suggesting that OPN from the host also contributes to tumor proliferation.

We independently validated the reduction in tumor cell proliferation using Ki67 immunohistochemistry (IHC) (Fig. 4M & N). As expected, the WTt-WTm samples contained more Ki67^+^ cells per view field than the knockout groups. To directly test the role of OPN in tumor cell proliferation, we performed in vitro proliferation assays. OPN-knockout CT26 and MC38-met cells exhibited a significant decrease in proliferation compared to scramble controls (Fig. 4O). Treating parental CT26 and MC38-met cells with αOPN antibody phenocopied the knockout effect, confirming that proliferation is OPN-dependent (Fig. 4P). To explore the underlying mechanism of OPN-driven tumor cell proliferation, we calculated a MAPK signaling score from relevant gene signature^45^ using genes from our dataset, revealing a decrease in MAPK activity in all knockout groups relative to WTt-WTm (Fig. 4Q). Western blot analysis showed significantly decreased MAPK kinase activity as determined by phosphorylation of MEK and ERK in OPN-knockout cell lines compared to scramble controls, consistent with known OPN signaling pathways (Fig. 4R-T)^46^. Furthermore, OPN promotes myeloid proliferation (Fig. SA-C), but via a MAPK-independent signaling pathway (Fig. S5A-F).

Notably, MAPK activity has been reported to activate PGE2 production in cancer cells, which can suppress monocytes and T cells; however, in our model, knocking out OPN from MC38met tumor cells did not increase PGE2 production (Fig. S5G &H)^47,48^. Taken together, these results demonstrate that autocrine OPN signaling in tumor cells drives proliferation via the MEK/ERK pathway.

### Tumor-and host-derived OPN regulate monocyte differentiation and macrophage polarization

One of the most striking differences in cell-type composition across the experimental four groups was a relative increase in monocytes in OPN-deficient hosts (WTt-KOm and KOt-KOm; Fig. 2E). Comparison of monocyte and macrophage proportions revealed that M2-like macrophages dominated in WTt-WTm tumors (Fig. 5A), whereas KOt-WTm tumors were enriched for M1-like macrophages. In OPN-deficient hosts (WTt-KOm and KOt-KOm), monocytes comprised most of this myeloid compartment (Fig. 5A-C). Differential expression analysis highlighted distinct transcriptional programs across the three clusters. M1-like macrophages expressed high levels antigen-presentation transcripts (*H2-Eb1*, *H2-Ab1*, *H2-Aa*, *Cd74*, and *H2-Dmb1*) (Fig. 5B). M2-like macrophages expressed complement components (*C1qa*, *C1qc*, *C1qb*) along with *Pf4* and *Mrc1* (CD206) (Fig. 5B), suggesting substantial overlap with previously described SPP1^hi^ TAMs that promote liver metastasis. Monocytes were characterized by high expressed of *Plac8*, *Mx1*, *Ifit3/b*, *Cxcl10*, and *Srgn*, indicative of an active, interferon-responsive state (Fig. 5B)^47^.

**Figure 5.**
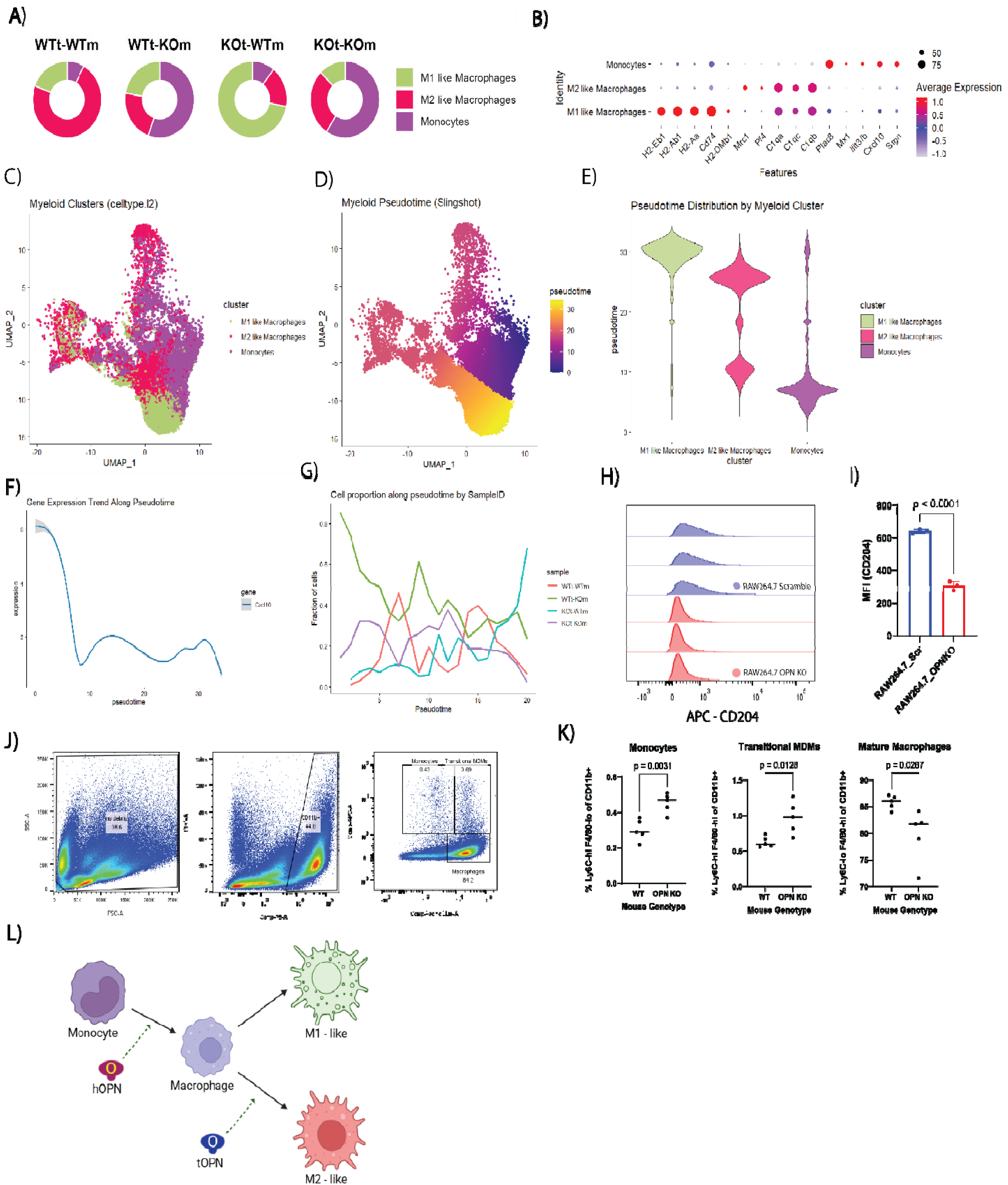
Host OPN drives monocyte-to-macrophage maturation, while tumor-derived OPN polarizes macrophages towards a pro-tumor M2 phenotype. (A) Donut plots showing proportions of M1-like macrophages, M2-like macrophages, and monocytes within tumor nodules across the four experimental groups. (B) Dot plot of differential gene expression between the three myeloid subtypes, sorted by log2FC. (C) UMAP of the indicated myeloid populations (D-E) Pseudotime analysis of myeloid clusters on UMAP (D) and corresponding violin plots of pseudotime distributions (E) (F-G) *Cxcl10* expression across pseudotime in myeloid cells (F) and proportion of myeloid cells versus pseudotime, with lines representing each experimental group. Proportion of myeloid cells versus pseudotime. Each experimental group has its own line (G). (H-I) Flow cytometry of CD204 on the RAW264.7 cell lines (scramble and OPN KO; n = 3 biological replicates) (J-K) Flow cytometry of monocytes, transitional monocyte-derived macrophages, and F4/80-hi mature macrophages in the peritoneal exudates of WT and OPN-KO mice. Gating scheme (J) and quantification (K; n = 5 biological replicates) L) Model summarizing host-and tumor-derived OPN regulation of monocyte differentiation and polarization (created with BioRender.com).

UMAP projection of these clusters and pseudotime analysis, with monocytes as the root cluster, revealed a transcriptional continuum from monocytes to macrophages (Fig. 5C-E). Most monocytes were positioned early in pseudotime, M2-like macrophages occupied intermediate states, and M1-like macrophages resided at the distal end, reflecting substantial transcriptional divergence (Fig. 5E). Consistent with this, *Cxcl10*, a master chemokine for recruiting CXCR3^+^ T and NK cells, was highest in early pseudotime (Fig. 5F), and the WTt-KOm and KOt-KOm groups contained the most cells in this region (Fig. 5G). These data suggest that in OPN-deficient hosts, monocytes with immune-recruiting potential dominate the myeloid compartment, fitting within the broader context of a recent studies showing monocyte populations capable of cross-presenting tumor antigens to T cells and facilitating anti-tumor immunity^47^.

To directly examine the role of OPN in macrophage polarization, we knocked out *Spp1* in RAW264.7 macrophages. These already differentiated cells model mature macrophage behavior in the TME rather than monocyte differentiation dynamics. Flow cytometry revealed that OPN-deficient RAW264.7 cells exhibited decreased CD204 (Macrophage Scavenger Receptor 1) compared to scramble controls (Fig. 5H & I, S5J), consistent with a reduction in M2-like, pro-tumor phenotype and with previous reports^49,50^. These results highlight that OPN acts in the TME in a paracrine and autocrine fashion to sustain alternative macrophage activation.

The expansion of monocytes in tumors from OPN-deficient hosts – but not in KOt-WTm tumors – suggested that host OPN regulates monocyte-to-macrophage differentiation prior to entry into the TME. To test this, we analyzed peritoneal exudate cells from tumor-free WT and OPN-KO mice, where monocyte recruitment is ongoing (Fig. 5L) ^51^. Flow cytometry revealed increased Ly6C^hi^, F4/80^hi^ and Ly6C-hi, F4/80-lo populations in OPN-KO mice relative to WT (Fig. 5J & K), indicating impaired differentiation and thus accumulation of monocytes in the absence of host OPN. These data support a model in which host-derived OPN licenses monocyte differentiation into macrophages, while tumor-derived OPN in the TME alternatively activates macrophages towards an M2-like, pro-tumor state (Fig. 5L).

### OPN suppresses T cells in the tumor microenvironment

T cells lie at the center of anti-tumor immunity. We previously demonstrated that OPN suppresses T cell proliferation and activation in colon tumors in vivo, focusing on *IRF8*-dependent myeloid mechanisms and global antibody-mediated OPN blockade^20,22,52^. Here, we extended these findings to dissect the compartment-specific roles of OPN in regulating T cell immunity. All three knockout groups (WTt-KOm, KOt-WTm, and KOt-KOm) exhibited increased T cell infiltration compared to WTt-WTm tumors (Fig. 6A). It was recently reported that SPP1 deficiency in myeloid cells upregulated CXCL9 and CXCL10 expression via the cGAS-STING pathway, which may explain the poor infiltration of T cells in OPN-competent TMEs^53^. The increase in T cells in the OPN-knockout groups was driven by niche-specific enrichment of T cells. Anti-tumor niches T6 and T7 – previously identified in OPN-deficient hosts and KOt-WTm tumors, respectively – contained the highest proportions of T cells among all niches (Fig. 6B). Visual inspection of representative FOVs confirmed that T6 was enriched for T cells, monocytes, and DCs, whereas T7 was enriched for T cells, M1-like macrophages, and DCs (Fig. 6C & D). Co-localization analysis revealed close spatial association between T cells and DCs in T7 (Fig. 6E).

**Figure 6.**
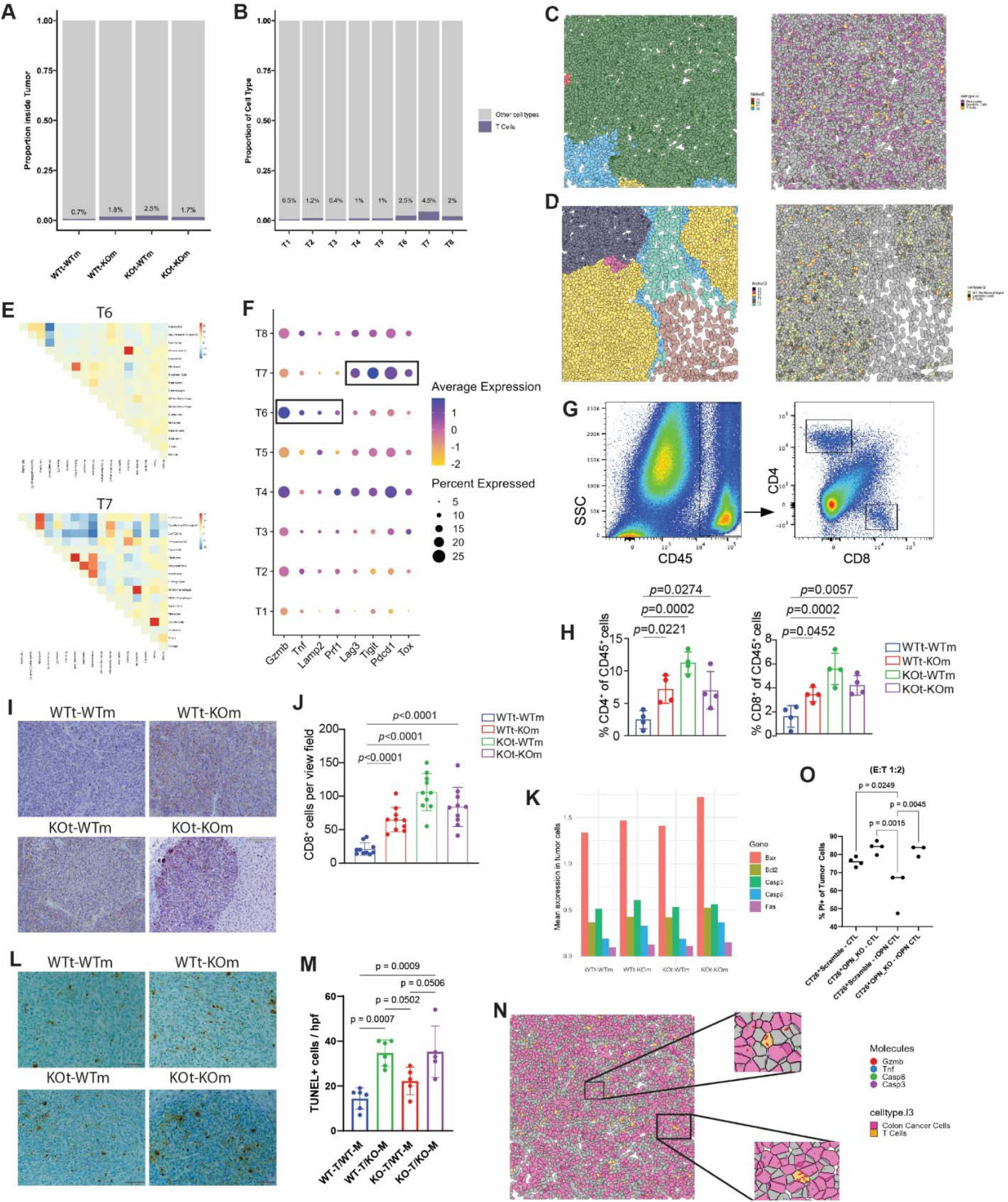
Tumor-and host-derived OPN suppress T cells in the tumor microenvironment. (A-B) Bar plots showing proportion of T cells inside tumor niches across samples (A) and niches (B). (C-D) Representative FOVs of niches T6 (C) and T7 (D) colored by niche (left) and niche-defining immune cells (right). (E) Co-localization analysis of cells within niche T6 (top) and T7 (bottom) (F) Dot plot of cytotoxicity-and exhaustion-associated markers in T cells across the tumor niches; boxes indicate selected features for emphasis. (G-H) Flow cytometry of T cells from liver metastasis tumor digests, including the gating strategy (G) and quantification (H; n = 4-5 biological replicates) (I-J) Representative images (I) and quantification (J) of CD8 IHC staining across the four experimental groups (n = 10 high-powered fields per group). (K) Bar plots showing mean expression of apoptosis-related genes across the four sample groups from the CosMx dataset. (L-M) Representative images (L) and quantification (M) of TUNEL IHC staining of the four experimental groups (n = 10 high-powered fields per group) (N) Representative FOV from (C) colored by tumor cells, T cells, and indicated transcripts. Insets highlight actively cytotoxic T cells expressing *Gzmb* co-localized with tumor cells expressing apoptosis-related transcripts. (O) 2/20 CTLs were pretreated with recombinant OPN (1 µg/mL, 3 days) or left untreated, then co-cultured with CT26 Scramble or OPN_KO cells (1E5) at an E:T ratio of 1:2. After 24 h, adherent and floating cells were collected, stained with CD8a and PI, and analyzed by flow cytometry to quantify tumor cell apoptosis.

Because the greatest tumor suppression occurred in the KOt-KOm group (Fig. 1K), which was preferentially enriched for niche T6 rather than T7, we sought to determine whether the functional state of T cells was different between these two niches. T cells in niche T6 expressed higher levels of cytotoxicity-related genes (*Gzmb, Tnf, Lamp2,* and *Prf1*), whereas T cells in niche T7 exhibited elevated expression of exhaustion-associated markers (*Lag3, Tigit, Pdcd1, Tox)* (Fig. 6F). We validated the increased infiltration of T cells in OPN-deficient tumors using flow cytometry and IHC. Consistent with the transcriptomics data, all three knockout groups exhibited increased T cell abundance relative to WTt-WTm tumors (Fig. 6G-J).

Given the heightened cytotoxic T cell signature in T6, we next examined whether tumor cells in OPN-deficient hosts exhibited evidence of increased susceptibility to immune-mediated killing. Although the CosMx panel did not assay active apoptosis, tumors from WTt-KOm and KOt-KOm groups showed increased expression of apoptosis-related genes (*Bax*, *Bcl2*, *Casp3*, *Casp8*, and *Fas*), particularly *Casp8* (Fig. 6K). While elevated expression of these genes does not directly indicate apoptosis, it suggests cellular stress and a state primed for cytotoxic attack. Consistent with this interpretation, TUNEL staining revealed significantly increased apoptosis in tumors from OPN-deficient hosts compared to WTt-WTm and KOt-WTm tumors (Fig. 6L).

Supporting active T cell-mediated killing, we identified regions within niche T6 with *Gzmb*-hi T cells near tumor cells expressing apoptosis-associated transcripts, providing spatial evidence of active cytotoxic interactions (Fig. 6N). To directly test whether OPN suppresses T cell cytotoxic function, we performed co-culture experiments using gp70-specific 2/20 CTLs and gp70-expressing CT26 tumor cells^54,55^. CTLs were more effective at killing CT26-OPN KO cells compared to scramble controls (Fig. 6O, lanes 1-2). Pre-treatment of CTLs with recombinant OPN significantly reduced their ability to kill CT26-Scramble cells (Fig 6O, lanes 1 & 3).

Importantly, removal of OPN-mediated suppression by culturing OPN-stimulated CTLs with CT26-OPN KO cells rapidly restored cytotoxic activity (Fig 6O, lane 4).

Together, these data demonstrate that OPN-deficient hosts are enriched for a tumor niche containing highly cytotoxic, non-exhausted T cells capable of effectively suppressing tumor progression. While loss of tumor-derived OPN increases T cell infiltration, these cells display a more exhausted phenotype. Thus, host-and tumor-derived OPN cooperate to suppress T cell infiltration, function, and cytotoxicity in the TME.

### Host OPN deficiency unlocks an interferon response niche driving anti-tumor immunity

After observing that T cells in niche T6 displayed an activated state, we sought to characterize the broader immune landscape of niche T6. Niche T6 was predominantly enriched in host-OPN deficient tumors (WTt-KOm and KOt-KOm), whereas T7 was meaningfully observed only in the KOt-WTm group (Fig. 7A). Niche T6 was characterized by a high abundance of monocytes and IFN-response tumor cells, while T7 was enriched in M1-like macrophages (Fig. 7B). Differential expression analysis revealed that T6 was enriched for genes involved in immune recruitment and type I interferon signaling (*Cxcl10*, *Plac8*, *Ifit3b*, *Ifit1*, *Oas3*) (Fig. 7C). In contrast, T7 was defined by antigen presentation programs (*H2-Ab1*, *Cd74*, *H2-Aa*, *H2-Eb1*), consistent with prior reports linking IFN phenotype and CD74 expression to ICI response in CRC^56^.

**Figure 7.**
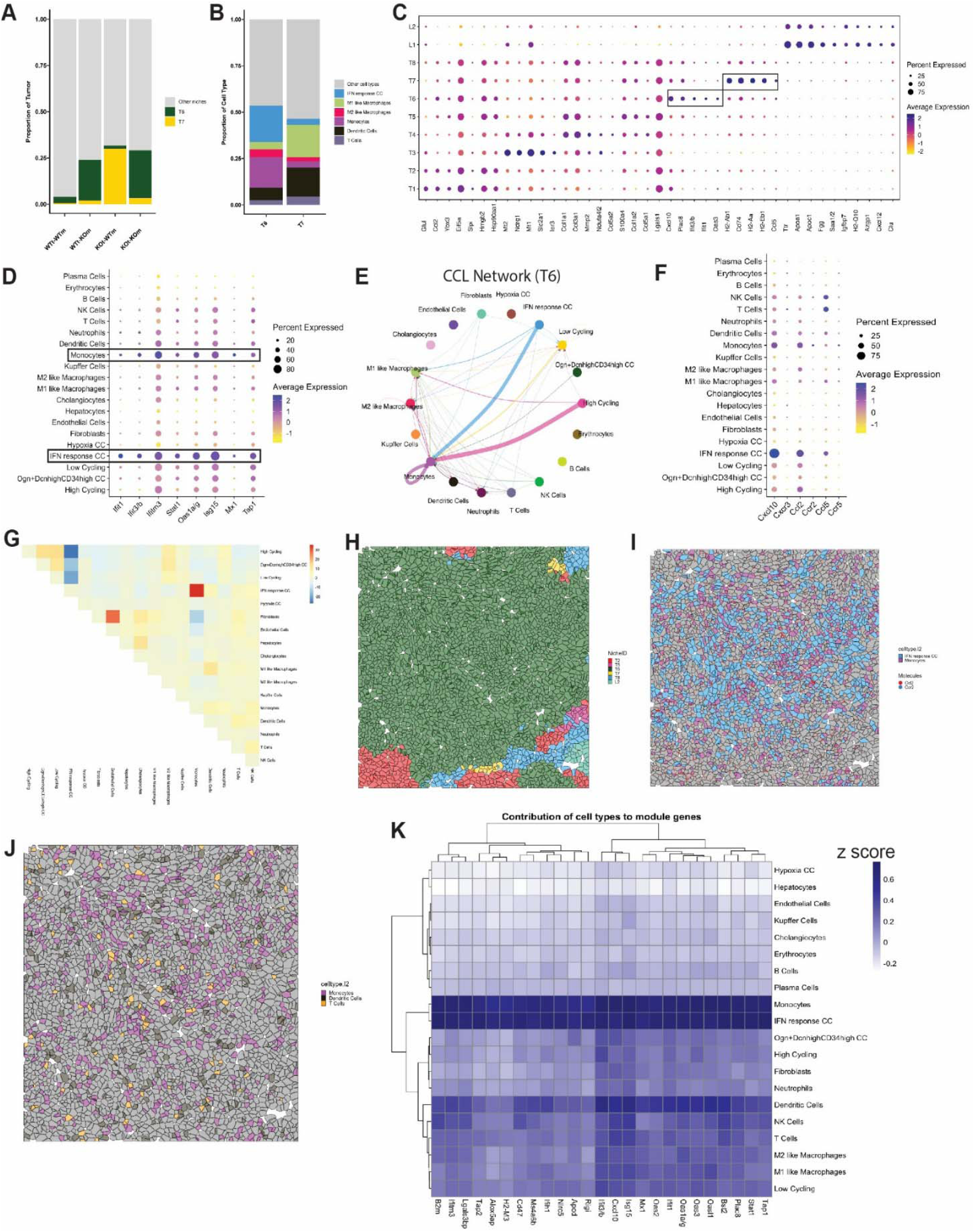
IFN response programs define an anti-tumor immunity niche in OPN-deficient hosts (A-B) Bar plots showing proportion of niches T6 and T7 across samples (A) and the cellular composition of these niches (B) (C) Dot plot of differentially expressed genes across tumor niches, sorted by log2 fold change; boxes indicate selected features for emphasis (D) Dot plot of IFN response program genes across cell types; boxes indicate cell types with high expression of this program (E) CellChat analysis of the CCL network (net aggregate) from a representative niche T6 field of view (FOV); connection thickness indicates total interaction strength. (F) Dot plot of selected chemokines and their cognate receptors across cell types (G) Co-localization heatmap of cell types in niche T6 (H-J) Representative FOV of niche T6 colored by niche (H), monocytes and IFN response cancer cells with indicated chemokine and receptor (I), and selected immune cell populations (J) (K) Estimated contribution of each cell type in each gene of the Isg15_Oas1a.g_25 module.

Given that the globally OPN-deficient TME (KOt-KOm) most closely models the likely effect of OPN-blockade immunotherapy, we investigated whether host OPN deficiency promotes a distinct anti-tumor niche not seen elsewhere. We found that monocytes and IFN-response cancer cells were the primary contributors to the IFN response program in T6 (Fig. 7D), suggesting that monocyte recruitment and local IFN signaling are key features of this niche.

Monocytes comprised a significant fraction of T6, consistent with active recruitment. CellChat analysis of the CCL signaling network indicated that IFN response cancer cells, high cycling cancer cells, and monocytes collectively signaled to recruit monocytes (Fig. 7E), corroborated by high expression of *Ccl2* and *Ccl5* in these cell types (Fig. 7F). Spatial analysis further confirmed co-localization of monocytes and IFN-response tumor cells within T6, which we verified through visual inspection of a representative T6 FOV (Fig. 7G-I, S&). Re-coloring the same FOV for adaptive immune cells showed that T cells frequently contacted both DCs and monocytes, suggesting that these innate populations may shape T cell activity within the T6 niche.

To further assess whether transcriptional programs mirrored the spatial organization of cells, we conducted a spatially colocalized gene module (SCGM) analysis using cell centroids as transcript locations. We identified 37 gene modules with high spatial co-localization, including a 25-gene module representing a type I interferon response program (Fig 7K, S8A). Consistent with our other analyses, monocytes and IFN-response tumor cells were the dominant contributors to this module (Fig. 7K), supporting the conclusion that T6 is a monocyte-drive, IFN-response niche.

### OPN blockade immunotherapy suppresses CRC liver metastasis and enhances T cell infiltration

Guided by our findings from genetic knockout experiments, we evaluated the therapeutic potential of OPN neutralization in vivo. Balb/c mice bearing CT26 liver metastasis were treated with an α-OPN monoclonal antibody (100D3). OPN blockade immunotherapy significantly reduced tumor burden in this syngeneic model (Fig. 8A). Flow cytometry and IHC revealed a corresponding increase in T cell infiltration following treatment (Fig. 8B-E).

**Figure 8.**
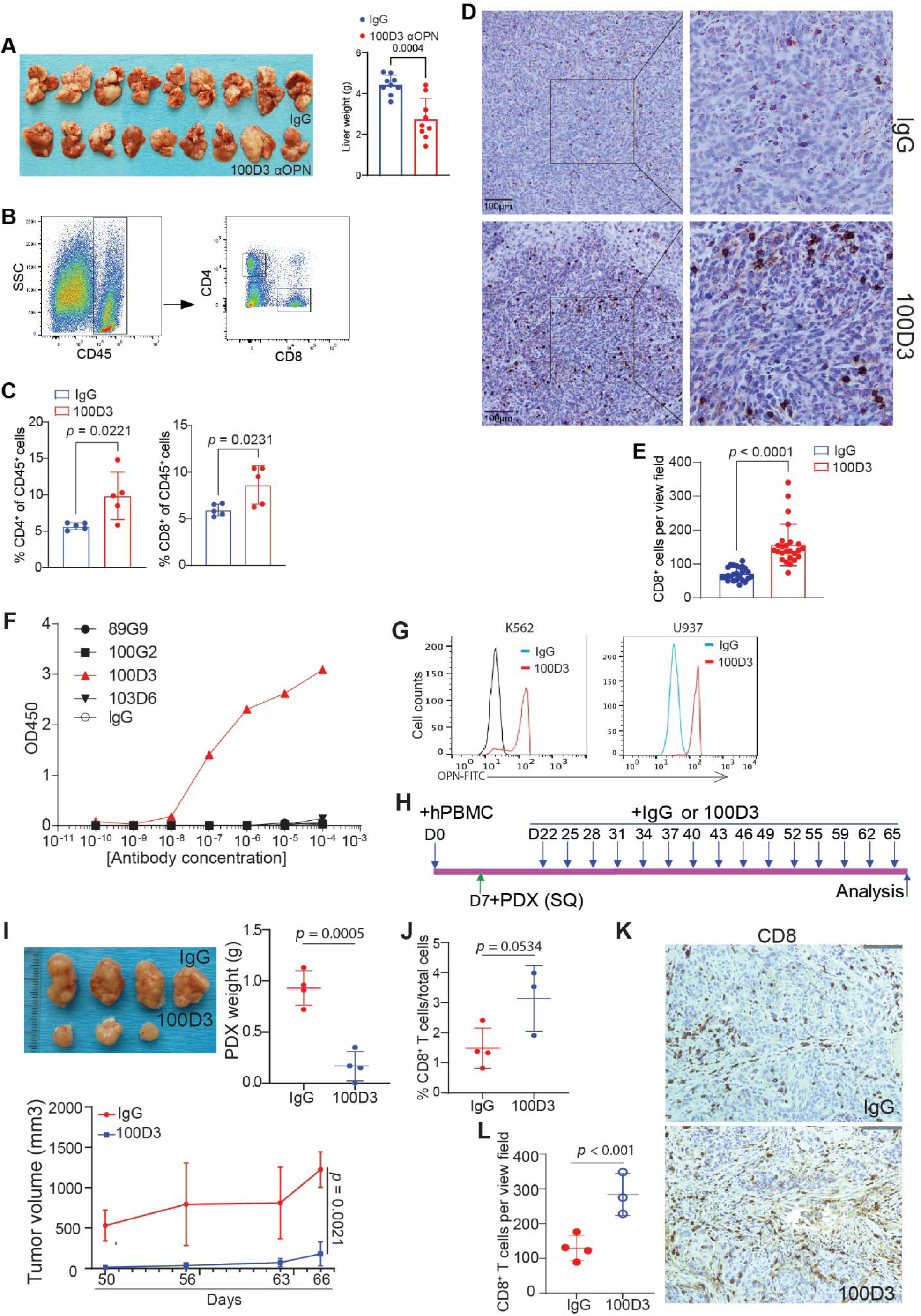
Osteopontin blockade reduces tumor burden and increases T cell infiltration in syngeneic and patient-derived xenograft mouse models. (A) CT26 cells were injected into spleens of BALB/c mice to establish liver metastasis. Mice were treated with IgG or anti-OPN mAb (clone 100D3) starting 5 days after tumor inoculation, every three days (q3d) for four doses. Livers were analyzed 17 days after tumor cell injection; images of tumor-bearing livers and quantification are shown (n = 9 biological replicates) (B-C) Flow cytometry analysis of tumors from (A) including the gating strategy (B) and quantification (C; n = 5 biological replicates). (D-E) CD8 IHC staining of tumors from (A). Representative images are shown (D) along with quantification (E; n = 25 view fields per group) (F) Indirect ELISA analysis of the indicated OPN mAbs in vitro. (G) Intracellular staining with 100D3 followed by flow cytometry analysis in human myeloid cells. (H) Schematic of 100D3 immunotherapy in patient-derived colon cancer liver metastases xenografts (PDXs) established in humanized NSG mice. (I) Representative PDX images (top left), tumor growth kinetics (bottom left) and tumor weight at the experimental end point (top right; n = 4 biological replicates) (J) Flow cytometry quantification of tumor-infiltrating CD8+ T cells in PDX in humanized mice at the endpoint (n = 3-4 biological replicates). (K-L) IHC staining of PDX tumors using a human-CD8α-specific antibody at the endpoint. Representative images are shown (K) and the results were quantified (L); brown signal indicates CD8 staining (n = 3-4 biological replicates).

To determine whether these effects translated to human CRCLM, we employed a human colon cancer patient liver metastases-derived xenograft (PDX) model in humanized mice. We first confirmed that our clone 100D3 binds human OPN protein in vitro (Fig. 8F) and intracellularly in myeloid cells (Fig. 8G). Mice bearing subcutaneous PDX tumors were treated with α-OPN or IgG every three days for 15 cycles (Fig. 8H). Tumors from the treatment group were significantly smaller, including one complete remission (Fig. 8I). Analysis by flow cytometry and IHC again demonstrated enhanced T cell infiltration in the treatment group (Fig. 8J-L).

Together, these results indicate that therapeutic OPN blockade suppresses tumor growth and promotes T cell infiltration in both syngeneic and humanized CRCLM models, consistent with our genetic studies.

## Discussion

In this study, we demonstrated that Osteopontin (OPN) promotes colorectal cancer liver metastasis through compartment-specific mechanisms. In our models, tumor-derived OPN drives tumor proliferation via the MEK/ERK pathway. Host OPN licenses monocyte differentiation into macrophage, whereas OPN in the tumor microenvironment (TME) promotes polarization to an M2-like, pro-tumor phenotype. Both tumor-and host-derived OPN contribute to the suppression of T cells in the TME. In OPN-deficient hosts, T cells were more cytotoxic and less exhausted than in OPN-deficient tumors, contributing to an IFN-response-program-driven niche that promotes anti-tumor immunity.

Our findings unify several observations from prior studies while adding critical single-cell and spatial resolution analysis. Previous work implicating SPP1 in CRC frequently relied on bulk expression analysis, which cannot resolve compartment-specific roles. At the single-cell level, a recent study showed that SPP1 blockade reduced F4/80^+^ TAMs in tumors – a phenomenon we also observe – but we additionally find that this reduction is accompanied by an increase in monocytes, highlighting a previously underappreciated shift in the myeloid compartment^42^.

Studies in melanoma have shown that inflammatory monocytes dominate treatment-sensitive tumors, whereas TAMs predominate in resistant tumors^47^. Given that these inflammatory monocytes are enriched for IFN-response program genes, the monocytes observed in niche T6 in our OPN-deficient hosts may serve an analogous anti-tumor function, suggesting that targeting SPP1 can reverse TAM-skewing that contributes to treatment resistance. The limited 1,000 gene panel used in CosMx SMI precludes direct mapping of previously described inflammatory monocyte signatures, but our findings are consistent with their functional role. As for the OPN-licensed differentiation of monocytes into macrophages, it remains to be studied whether this is due to intracellular OPN expression in myeloid cells or the result of extracellular OPN engaging surface receptors.

Recent work has identified an interferon-high immunophenotype as predictive of response to immune checkpoint inhibition (ICI) ICI in CRC^56^. Patients with this immunophenotype at baseline exhibit higher responsiveness to PD-1 blockade. In our study, the KOt-KOm group – which models global OPN blockade – shows enrichment for an IFN-response-drive niche (T6), suggesting that OPN blockade may unlock similar immune-activated states, which has been shown in mouse models to enhance responsiveness to ICI^42^.

Previous reports describe tumor-derived prostaglandin E2 (PGE2) as a mediator of T cell suppression in melanoma and other cancers^47,48^. We found that MC38met-OPN_KO cells exhibit increased PGE2 expression in vitro, indicating that the immune effects in our model are PGE2-independent and specifically attributable to OPN. Investigating dual targeting of PGE2 and OPN to reactivate T cells in the TME remains an intriguing area for future study.

Our results further delineate how tumor-and host-derived OPN cooperate to suppress T cells in a niche-dependent manner. We previously identified an OPN-CD44 immune checkpoint controlling CD8 T cell activation, and our current work is consistent with additional macrophage-mediated suppression mechanisms^20^. Literature shows that SPP1+ TAMs or FasL+CD11b+F4/80+ monocyte derived macrophages cause exhaustion or apoptosis of tumor-antigen specific T cells, leading to tumor progression^34,35^. Here, we show that targeting host-derived OPN impairs monocyte-to-macrophage differentiation, limiting the emergence of these pro-tumor populations. If the differentiation to macrophages occurs, OPN in the TME promotes M2-like polarization. Recent work identifies PI3K/AKT-mediated CSF1 signaling as a dominant pathway for this polarization^57^. Although CosMx SMI provides critical spatial information, it yields lower per-cell transcriptomic depth than current scRNAseq technologies. To understand how host-and tumor-derived OPN affect T cell differentiation and exhaustion states in CRCLM, future studies may consider the use of scRNAseq.

Safety of potential therapeutic targets is a critical aspect of preclinical and clinical drug development. One clinical trial has evaluated OPN-blocking monoclonal antibodies in humans for the treatment of rheumatoid arthritis (RA)^58^. While OPN blockade was not efficacious in RA, the favorable safety profile observed in the trial suggests that OPN may represent a viable therapeutic target in CRCLM.

Collectively, our study provides mechanistic evidence that OPN cooperates in a compartment-specific manner to support liver metastasis. These results support the rationale for global OPN targeting as a therapeutic strategy to reduce tumor proliferation and relieve multiple layers of host-OPN driven immune suppression that leads to metastatic progression in colorectal cancer.

## Methods

### Tumor cell lines

Mouse colon tumor cell line CT26, mouse leukemia cell line RAW264.7, and human leukemia cell lines K562 and U937 were obtained from American Type Culture Collection (ATCC) (Manassas, VA). MC38-met is a metastatic mouse colon tumor cell subline derived from the parent MC38 cell line^59^. J774M cell subline was derived from the parent J774A.1 cell line from ATCC^60^. ATCC characterized these cells by morphology, immunology, DNA fingerprint, and cytogenetics. Cells were negative for mycoplasma at the time of experiments.

### Mice

BALB/c and C57BL/6 mice were obtained from Charles River NCI Frederick (Frederick, MD). NSG (NOD.Cg-*Prkdc^scid^ Il2rg^tm1Wjl^*/SzJ) mice were obtained from the Jackson laboratory (Bar Habor ME).

### Generation of OPN knock out cell lines

To generate *Spp1* knock out cell lines, HEK293FT cells were co-transfected with psPAX2 (Cat#12260, Addgene, Watertown, MA, USA), pCMV-VSV-G (Cat#8454, Addgene), and the scramble or *Spp1*-sgRNA-coding plentiCRISPRv2 (Genscript, Piscataway, NJ, USA) plasmids by Lipofectamine 2000 (Life Technologies, Carlsbad, CA, USA). *Spp1* and scramble sgRNA sequences are 5′-AAGGTGAAAGTGACTGATTC-3′ and 5′-CTCGTATCTTTTCCCACGGC-3′. After 48-72 hours, the virus particles were harvested to transduce CT26, MC38-met, RAW264.7, and J774M cells. The transduced cells were then cultured in the presence of puromycin (5 µg/mL) to select stably transduced cells. Cell phenotype was confirmed by analysis of OPN protein level in the cell culture supernatant.

### Colon tumor experimental liver metastasis syngeneic mouse model

The experimental mouse colon tumor liver metastasis syngeneic mouse model was established according to the published procedures^43^. Briefly, MC38-met or CT26 cells (3×10^5^ cells in 50 μl PBS) were surgically injected into the spleen of mice, followed by splenectomy. Use of mice is approved by Augusta University IACUC (protocol # 2008-0162).

### Human colon cancer liver metastases-derived PDX humanized mouse model

The human colon cancer patient liver metastases-derived PDX (TM00170) was established from a woman with colon cancer and obtained from the Jackson Laboratory. Humanized NSG mice were generated as previously described ^61^. Briefly, PBMCs were collected from consented healthy female donors at Augusta Shepeard Community Blood Center. PBMCs (3×10^6^ cells/mouse) from was injected iv to female NSG mice. This PDX humanized mouse model was determined to be non-human subject research and approved for use by Augusta University IRB (Protocol # 1314554-18).

### CosMx™ spatial molecular imager (SMI) sample preparation

Liver tissues were fixed in 10% buffered formalin and embedded in paraffin. Tissue was sectioned at 4 µm onto Leica BOND Plus slides, baked at 37 °C overnight, and stored at 4 °C until ready for CosMx SMI preparation. To improve adherence, the slides were baked overnight at 60 °C. After baking, samples were prepared according to the protocol provided by Bruker Spatial. Briefly, slides were deparaffinized and prepared for antigen retrieval. Immediately following antigen retrieval, slides were rinsed in DEPC-treated water and washed for 3 minutes in 100% ethanol. Samples underwent mild digestion with proteinase K. After rinsing, slides were incubated with 0.001% fiducial solution and fixed with 10% NBF. Following fixation, samples were quenched and then blocked using 100 mM NHS-acetate in CosMx NHS-acetate buffer.

Mouse Universal Cell Characterization Panel probes were denatured at 95 °C for 2 minutes, then snap-cooled on ice. The ISH probe working mix was prepared using 1 nM ISH probes in DEPC-treated water with Buffer R and RNase Inhibitor provided by Bruker Spatial. Samples were incubated with the probe mix at 37 °C overnight using coverslips and a hybridization oven for 16 hours.

Slides were then washed and incubated with DAPI nuclear stain diluted in blocking buffer. After appropriate washes, slides were incubated for 1 hour with morphology antibodies against CD298/B2M, PanCK, and CD45. After three washes in 1X PBS the glass flow cell was applied to the slide and slides were loaded into the instrument. Upon completion of the preview scan, fixed-size fields of view (FOVs) measuring 0.5 mm × 0.5 mm were selected to define the regions to be imaged for transcript detection. Eight Z-stack images (0.8 µm step size) were acquired per FOV. Photo-cleavable linkers on the fluorophores of the reporter probes were released by UV illumination and washed with strip wash buffer. The fluidic and imaging procedures were repeated for the 16 reporter pools. Each round of reporter hybridization and imaging was repeated multiple times to increase RNA detection sensitivity. Cell morphology was imaged on-instrument prior to RNA readout by adding fluorophore-bound reporters to the flow cell and capturing eight Z-stack images in the following channels: 488 nm (CD298/B2M), 532 nm (PanCK), 594 nm (CD45), 647 nm, and 385 nm (DAPI).

### Single cell segmentation

Data was transferred in real-time to AtoMx SIP software. Post-run quality controls were performed to ensure each FOV had a high hybridization rate and accurate cell segmentation, resulting in reliable transcript-per-cell quantification. Cell segmentation was further improved using AtoMx SMI advanced settings, optimized to reflect the tissue composition. Slide 1, with 331 FOV shows a mean per transcript per cell of 277 genes with a total of 740,902 non-empty cells. Slide 2 with 213 FOV shows a mean per transcript per cell of 348 genes with a total of 438,449 non-empty cells.

### Bioinformatics Analysis of CosMx Spatial Transcriptomics Data

The Seurat object exported from AtoMx SIP analysis was used for subsequent analysis. Cells failed to pass the quality check in AtoMx, and cells with fewer than 50 counts were removed from the downstream analysis. After the first pass of analysis as described below, the FastReseg package was used to evaluate the segmentation quality. A small number of cells (less than 0.02% of total cells) were removed if significant actions were taken by the FastReseg algorithm. The updated count matrix was then used for the second round of analysis. The updated count matrix was normalized using SCTransform. Batch correction was performed using the harmony algorithm. Cell type annotation was performed using the Seurat reference mapping or label transfer method. An artificial reference dataset was generated by integrating a scRNA-seq dataset of subcutaneously implanted MC38 cells in syngeneic mice (ref) and the liver single-cell atlas dataset (ref). Differential expression analysis was performed using the Seurat FindMarker function. Geneset signature scores were calculated using Seurat AddModuleScore function. Genesets were downloaded from MSigDB database. The niche analysis was performed in AtoMx and the niche labels were included in the AtoMx export and used for plotting figures. All visualization plots were generated using built-in plotting functions in Seurat. For computational analyses, *Spp1* transcripts were retained in the *Spp1*-knockout samples to preserve the integrity of cell-type clustering and prevent artifactual splitting due to the absence of *Spp1* transcripts. For clarity in representative images, *Spp1* transcripts were removed from KO tumor cells and/or host cells to highlight the absence of functional OPN protein.

### Cell co-localization analysis

We analyzed how different annotated cell types are positioned relative to one another in CosMx spatial transcriptomics data using a distance-based neighborhood graph built from cell centroid coordinates (in microns). For each sample (and, if needed, each field of view [FOV]), we defined neighboring cells as those within a fixed radius (e.g., 40 µm). We converted these relationships into a binary adjacency matrix indicating spatial contact. For a given “index” cell type A, we counted neighboring cells of type B by summing adjacency connections from A cells to B cells. To test whether this proximity was higher than expected by chance while preserving tissue structure, we created a null distribution by shuffling cell-type labels among non-index cells, either across the sample or within FOVs to maintain local density and spatial organization. From this null, we obtained expected neighbor counts, standard deviations, and z-scores for each cell-type pair and used them to compute two-sided p-values with multiple-testing correction. For regional analysis, we applied this method to each FOV, calculating enrichment scores (e.g., log of observed/expected neighbors). This highlighted spatial “hotspots” in which specific interactions, such as T cells and Dendritic cells or monocytes and IFN-response CC proximity, are particularly enriched.

### ELISA assays

Cell culture supernatant was collected, cleared by centrifugation, and measured for OPN protein concentration using the mouse OPN ELISA kit (R and D System, Minneapolis, MN) according to the manufacturer’s instructions. For antibody binding affinity assay, recombinant OPN protein was coated onto 96-well tissue culture plate at 1 μg/mL overnight. The wells were washed, blocked, and then anti-OPN antibody was added to the coated wells at various dilutions. Peroxidase-AffiniPure goat anti-mouse IgG was added as secondary antibody to capture the anti-OPN antibody. Assays were performed using the ELISA kit (Biolegend, San Diego, CA, USA) according to the manufacturer’s instructions.

### Cellular proliferation assay

Cells were seeded in 96-well plates and measured for cellular proliferation using the CellTiter 96A Queous One Solution Cell Proliferation Assay Kit (Cat# G3582, Promega Corp, Madison, WI).

### Generation of OPN monoclonal antibodies

To generate anti-human OPN neutralization monoclonal antibodies, five BALB/c mice were immunized with 50 μg recombinant human OPN protein (R and D System), followed by boost with 25 μg human OPN protein once every 2 weeks for 3 times. Mouse sera were tested by indirect ELISA using recombinant human OPN protein and a His tagged irrelevant protein as antigens. Based on OD value >1.0 while dilution is ≥1:8,000), mouse M6766 was selected for cell fusion. Spleen B cells were then fused with SP2/0 myeloma cells. Parent hybridoma clones were selected based on ELISA screening and selected for subcloing. Twenty clones were screened by ELISA for binding affinity and for suppressing proliferation of human leukemia cell lines K562 and U937 in vitro. Clones 13D11, 16B3, 16G2, 22B3, and 24C-3 were selected as the final products. Clone 100D3 was generated as previously described^22^.

### Western blotting analysis

Cells were lysed in total protein lysis buffer (20 mM of HEPES, pH 7.4, 20 mM of NaCl, 10% glycerol, 1% TritonX100, Proteinase inhibitor cocktails), centrifuged to remove debris, and separated in a 4–20% gradient denaturing SDS-polyacrylamide gel (Biorad, Hercules, CA, USA), transferred to PXDF membrane, blocked in 5% milk in PBST, incubated with primary antibodies (anti-pMek1/2, Cat#562457,and anti-pErk1/2, cat#612359, BD Biosciences) overnight at 4°C, followed by 2nd antibody and detection using the ECL system. The membrane was stripped and re-probed with anti-β-actin antibody (Sigma-Aldrich, Cat# 9026) as previously described^62^.

### Human colon cancer scRNA-seq datasets

The published human colon cancer patient liver metastases scRNA-seq datasets were extracted from GEO database (accession # GSE164522)^63^. Human colon tumor scRNA-seq datasets were extracted from GEO database (accession # GSE178341)^64^. Human colon cancer scRNA-seq datasets of patients treated with pembrolizumab were extracted from GEO database (accession # GSE236581)^65^. The datasets were analyzed using Seurat v4.9.9 in R v4.4.3.

### Human colon cancer survival and progression analysis

The TCGA Pan-Cancer (PANCAN) dataset was accessed via the Xena Browser program, and columns for Sample ID, *SPP1* expression, cancer type, sample type, OS, OS time, and AJCC stage were selected. Samples missing necessary values were excluded, retaining only COAD and READ primary tumors. Data were analyzed in R v4.4.3 using the tidyverse, survival, and survminer packages. Samples were classified as *SPP1*-high or *SPP1*-low (50% cutoff) and compared using Kaplan-Meier survival analysis. Tumor sub-stages were collapsed into parent stages (I-IV) for one-way ANOVA of *SPP1* expression across stages.

### Immunohistochemistry

Tumor tissues were fixed in 10% formalin and embedded in paraffin. The tumor sections were then processed and treated with HIER antigen retrieval agent (Abcam, Cat# ab208572), washed, blocked with serum, and incubated with anti-mouse CD8 antibody (Cat# Ab209775, Abcam) for mouse tumor tissues and with anti-human CD8 (Dako) for human tumor tissues. The stained tumor sections were counterstained with hematoxylin (Richard-Allan Scientific, Kalamazoo, MI) and then incubated with mouse IHC polymer detection kit (catalog no. ab209101, Abcam) to detect the target protein.

### Flow cytometry analysis

Tumor tissues were collected from mice and digested with collagenase solution [1 mg/ml collagenase (Sigma-Aldrich), 0.1 mg/ml Hyaluronidase (Sigma-Aldrich), and DNase I at 30 U/mL]. The digested tumor tissue was passed through a 100 μM cell strainer (Corning, Corning, NY), treated with red cell lysis buffer, and washed with 1% FBS in PBS. The mouse tumor digests were stained with fluorescent dye-conjugated anti-mouse CD45, CD4, CD8, CD11b, F4-80, Ly6C, Ly6G mAbs (Biolegend, San Diego, CA). The human colon cancer PDX digests were stained with fluorescent dye-conjugated anti-human CD8 antibody (Biolegend). For intracellular staining, cells were fixed, permeabilized, and stained with anti-OPN antibody, followed by staining with anti-mouse IgG-FITC. Stained cells were analyzed on a BD LSRFortessa cell analyzer with BD Diva 8.01 (BD Biosciences) or a BD FACSCalibur Flow cytometer with Cell Quest Pro. All flow cytometry data was analyzed with FlowJo v10.6.0 (BD Biosciences).

### Statistical Analysis

All **s**tatistical analysis was carried out using GraphPad Prism 10. ANOVA and paired Student’s t-test were used to determine statistical significance. A *p*<0.05 is considered as statistically significant.

## Acknowledgements

We acknowledge the support and contribution of the Integrated Genomics Core Shared Resources at the Georgia Cancer Center, Augusta University (RRID: SCR_026483). We thank Donna Kumiski in the Electron Microscopy & Histology Core for assistance in histology and Kimya Jones in the Georgia Esoteric & Molecular (GEM) Laboratory for assistance in IHC.

## Supplemental Materials

Supplemental figures 1-8

## Author Contributions

Conceptualization (PC, YZ, JDK, PS, HS, KL), Methodology: (PC, YZ, HS, KL), Investigation: (PC, YZ, JDK, KF, ZT, DP, MZ, KC, PSR, PVS, HS, KL), Formal Analysis (PC, YZ, HS, JW, KL), Funding acquisition: (KL), Project administration (KL), Software (PC, HS), Visualization (PC, YZ, HS, KL), Writing – original draft: (PC, KL), Writing – review & editing: (PC, YZ, JDK, KF, ZT, DP, MZ, KC, PSR, PVS, JW, HS, KL)

## Data availability

The Spatial transcriptomics datasets are deposited in GEO database. Accession #GSE309155.

