## Supplemental Figures for "Distinct and cooperative roles of host and tumor Osteopontin in colorectal cancer liver metastasis"

**Supplemental Data**

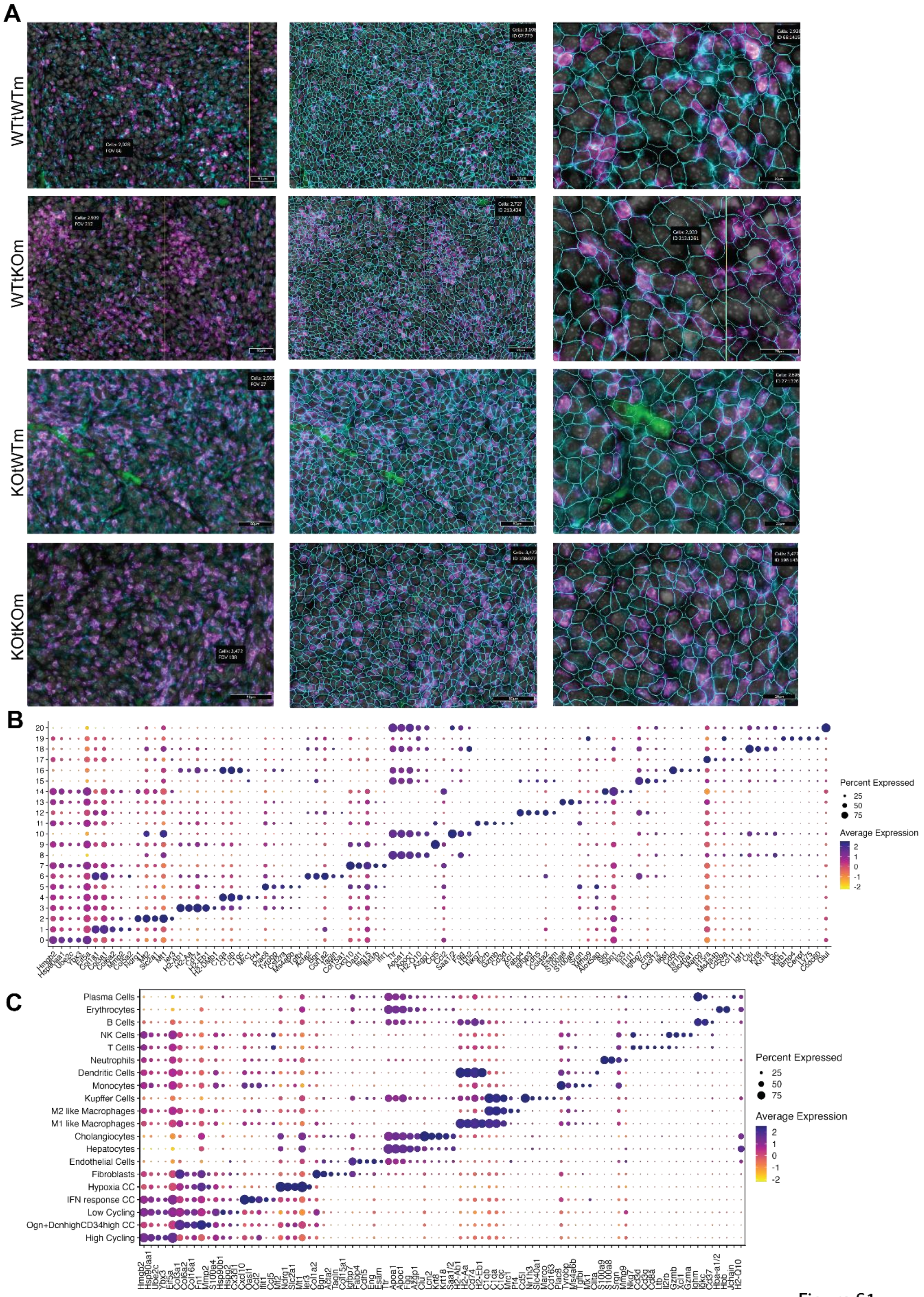


Figure S1

**Figure S1. Cell clustering and single cell segmentation**

A) Left panels: Sample data from Atomx SIP. Immunofluorescence composite image of the samples from the CosMx experiment in Fig 2A. Pan-CK (green), CD298/B2M (blue), CD45 (pink), DNA (grey). Middle panels: Images from left panel with cell segmentation borders overlayed. Right panels: magnified view of middle panels.
B) Differential expression dot plot of 21 unsupervised cell clusters, ranked by log2FC
C) Differential expression dot plot of annotated cell types, ranked by log2FC.

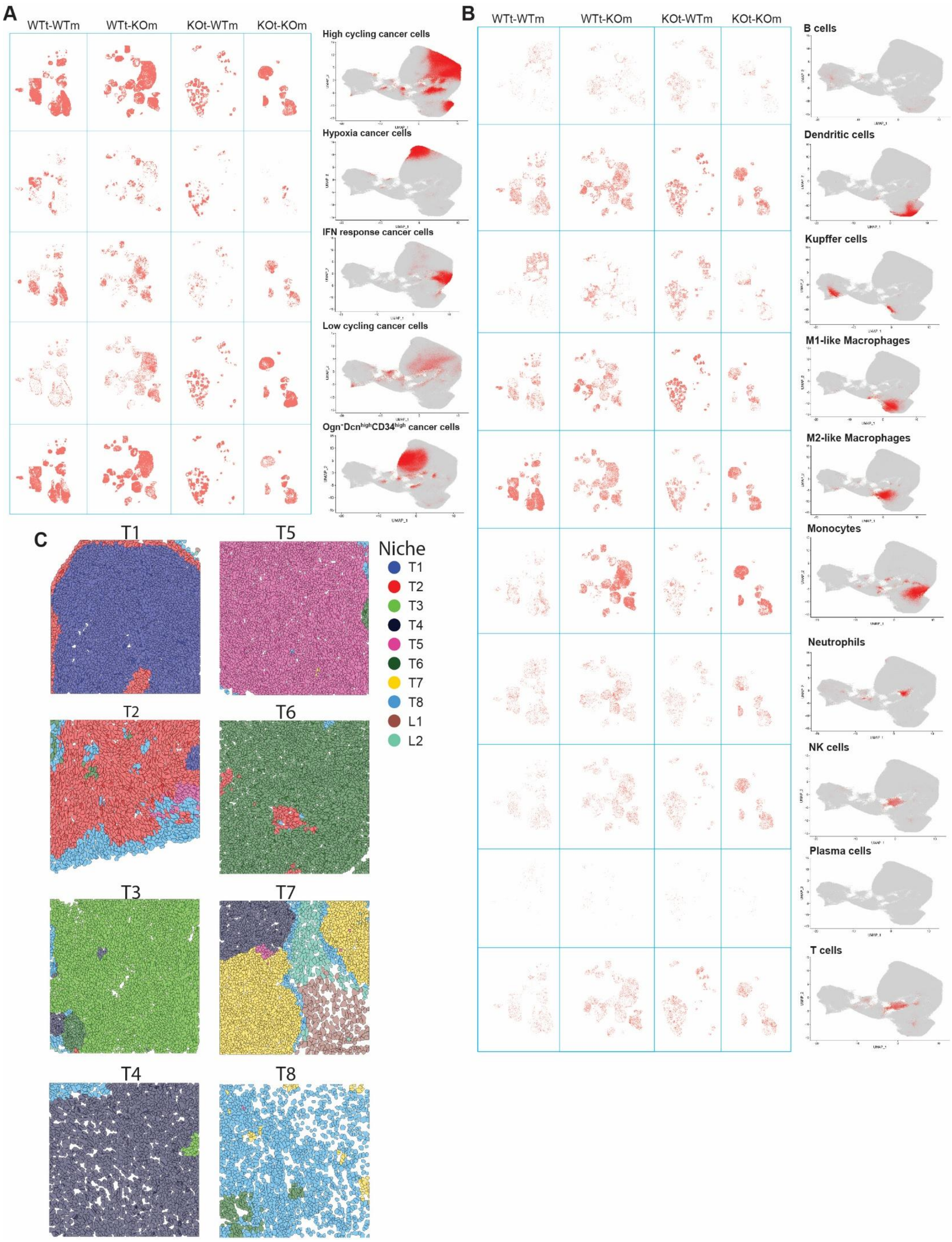
A
WTt-WTm
WTt-KOm
KOt-WTm
KOt-KOm
High cycling cancer cells
Hypoxia cancer cells
IFN response cancer cells
Low cycling cancer cells
Ogn⁻Dcn^high CD34^high cancer cells
B
WTt-WTm
WTt-KOm
KOt-WTm
KOt-KOm
B cells
Dendritic cells
Kupffer cells
M1-like Macrophages
M2-like Macrophages
Monocytes
Neutrophils
NK cells
Plasma cells
T cells
C
T1
T5
T2
T6
T3
T7
T4
T8
Niche
T1
T2
T3
T4
T5
T6
T7
T8
L1
L2

**Fig S2. Spatial and UMAP distribution of cell types**

(A-B) Spatial distribution maps and UMAPs with red dots indicating the spatial and UMAP positioning of tumor cell subtypes (A) and other cell types (B).

(C) Representative FOVs of the eight tumor niches from Fig 3F, colored by tumor niche

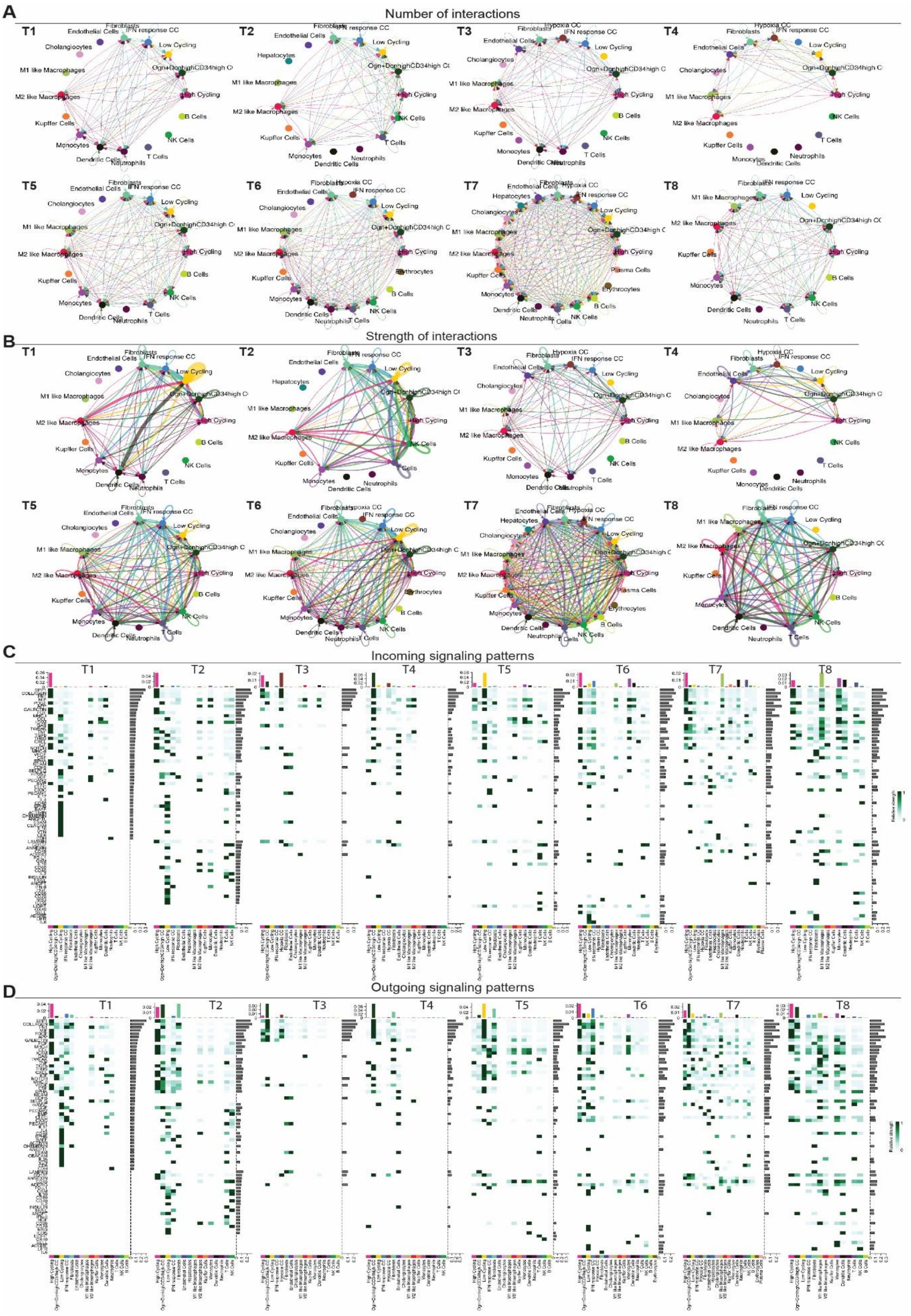
A
Number of interactions
T1
T2
T3
T4
T5
T6
T7
T8
Fibroblasts
IFN response CC
Hypoxia CC
Endothelial Cells
Cholangiocytes
Hepatocytes
Low Cycling
Ogn+DcnhighCD34high CC
M1 like Macrophages
M2 like Macrophages
High Cycling
Kupffer Cells
B Cells
NK Cells
Monocytes
T Cells
Dendritic Cells
Neutrophils
Erythrocytes
Plasma Cells
B
Strength of interactions
C
Incoming signaling patterns
D
Outgoing signaling patterns
Relative strength

**Figure S3. CellChat analysis results.**
(A-B) CellChat cell-cell communication analysis of representative Field of Views (FOV) of T1 to T8 niches based on known ligand–receptor pairs; Nodes represent cell types. Node size reflects the number (A) or total communication strength (B) of the cell population. Directed edges represent predicted signaling interactions, with arrows indicating the direction from ligand-expressing sender cells to receptor-expressing receiver cells. (C-D) Heatmaps show the incoming (C) and outgoing (D) signaling patterns of the representative FOVs in T1 to T8 niches. A gradient of white to dark green indicates low to high expression weight value in the heatmap.

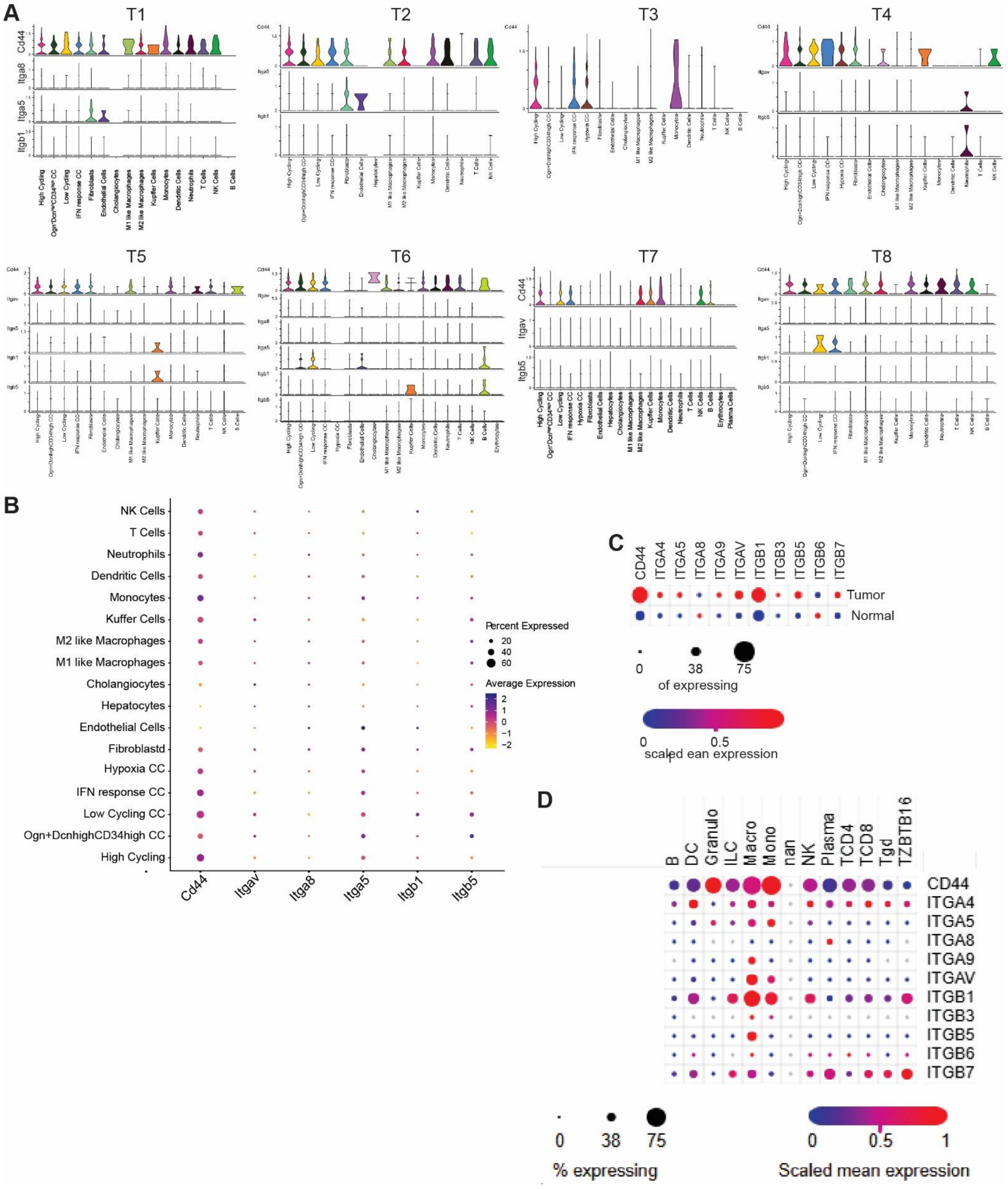
A
T1
T2
T3
T4
T5
T6
T7
T8
Cd44
Itga8
Itga5
Itgb1
Itgav
Itgb5
High Cycling
Low Cycling
IFN response CC
Hypoxia CC
Fibroblasts
Endothelial Cells
Hepatocytes
Cholangiocytes
M1 like Macrophages
M2 like Macrophages
Kupffer Cells
Monocytes
Dendritic Cells
Neutrophils
T Cells
NK Cells
B Cells
Erythrocytes
Plasma Cells
B
NK Cells
T Cells
Neutrophils
Dendritic Cells
Monocytes
Kuffer Cells
M2 like Macrophages
M1 like Macrophages
Cholangiocytes
Hepatocytes
Endothelial Cells
Fibroblastd
Hypoxia CC
IFN response CC
Low Cycling CC
Ogn+DcnhighCD34high CC
High Cycling
Cd44
Itgav
Itga8
Itga5
Itgb1
Itgb5
Percent Expressed
20
40
60
Average Expression
2
1
0
−1
−2
C
CD44
ITGA4
ITGA5
ITGA8
ITGA9
ITGAV
ITGB1
ITGB3
ITGB5
ITGB6
ITGB7
Tumor
Normal
0
38
75
of expressing
0
0.5
scaled ean expression
D
B
DC
Granulo
ILC
Macro
Mono
nan
NK
Plasma
TCD4
TCD8
Tgd
TZBTB16
CD44
ITGA4
ITGA5
ITGA8
ITGA9
ITGAV
ITGB1
ITGB3
ITGB5
ITGB6
ITGB7
0
38
75
% expressing
0
0.5
1
Scaled mean expression

**Figure S4**

(A) Violin plots of the expression of Spp1ligand and receptors of the SPP1 network in various cell types within T1 to T8 niches, respectively.

(B) The overall expression patterns of Spp1ligand and receptors in each cell types of the entire dataset

(C) Dot plot showing OPN expression profiles in human colon cancer using published human colon cancer scRNA-seq datasets.

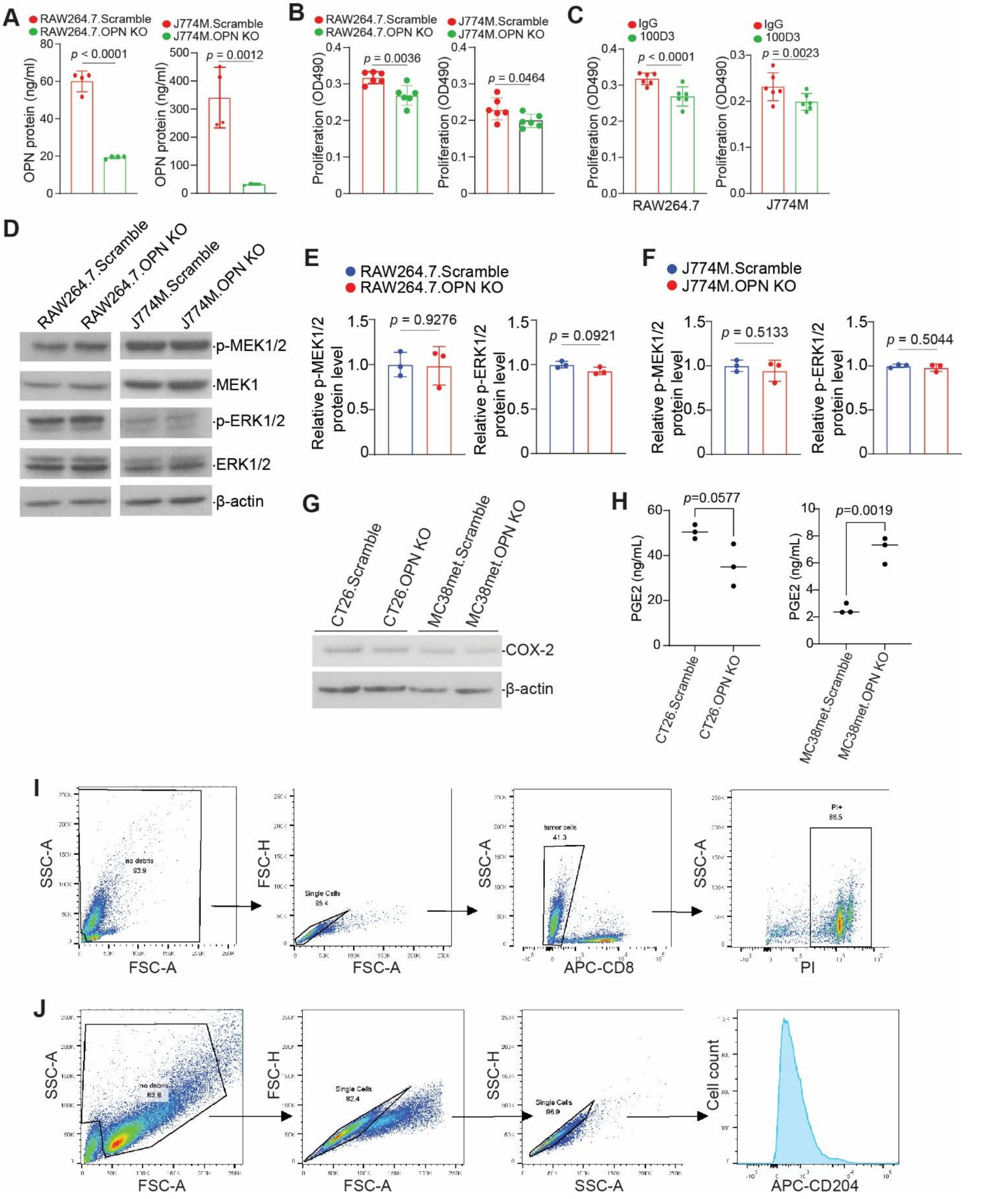

**Figure S5. Myeloid proliferation PGE2 production in tumor cells**

A) OPN concentration in cell culture supernatant of J774M Scramble, J774M OPN_KO, RAW264.7 Scramble, and RAW264.7 OPN_KO cell lines.
B) Proliferation assay of the cell lines from (A)
C) Proliferation assay of J774M and RAW264.7 treated with IgG or 100D3 (αOPN).

(D-F) Western blot analysis of MEK/ERK in the indicated cell lines. Myeloid proliferation is affected by OPN, but not through the MEK/ERK pathway as observed in tumor cells.

(G) Western blotting of Cox2 in the indicated cell sublines.

(H) ELISA analysis of PGE2 level in culture supernatants of the indicated cell sublines.

(I) Gating scheme for co-culture killing assay experiment in Figure 6O

(J) Gating scheme for RAW264.7 cell line staining for CD204 experiment in Figure 5H-I.

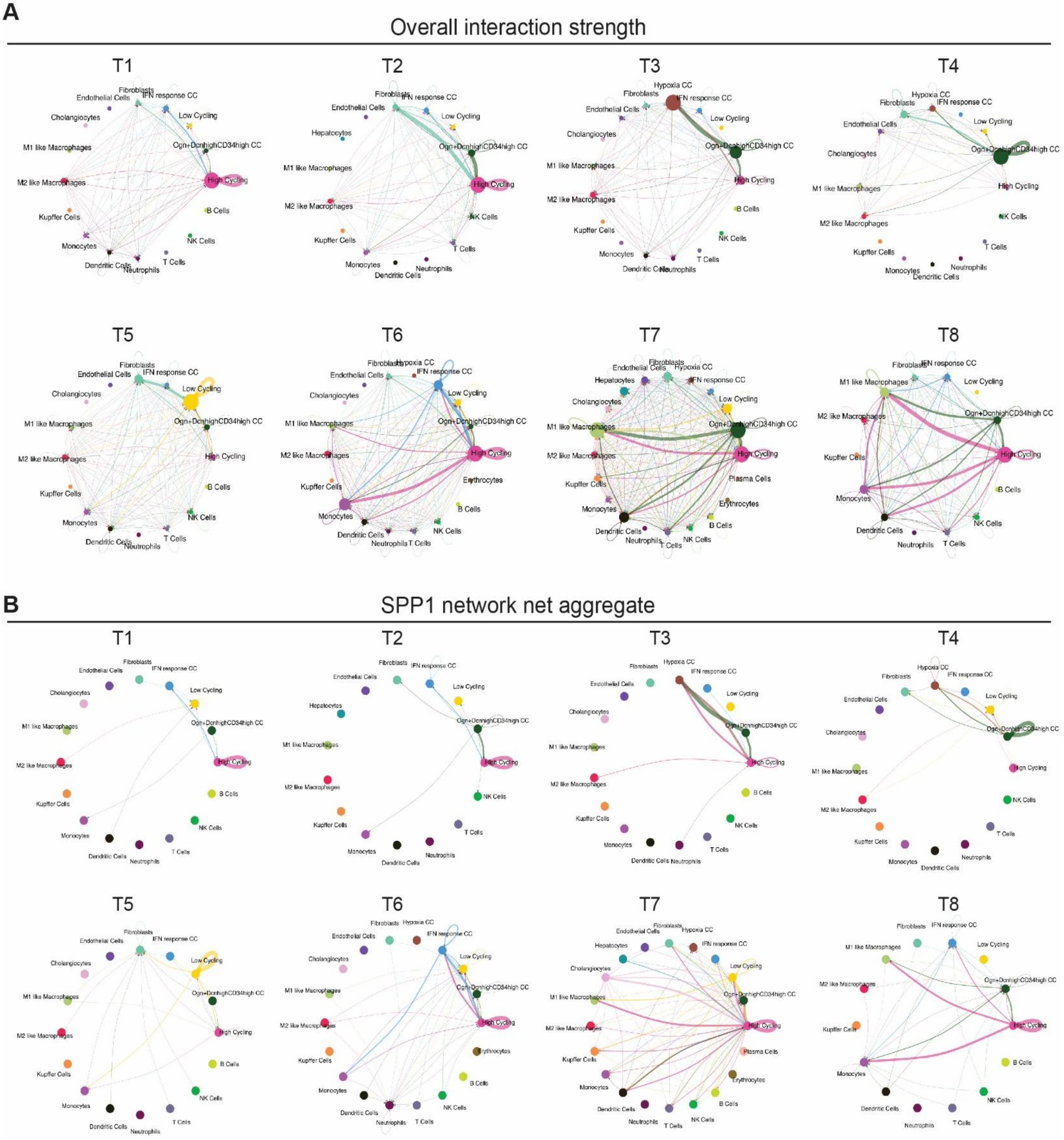


**Figure S6.** CellChat data

A) Cell chat overall interaction strength calculated on representative FOVs of a single niche
B) Cell chat net aggregate of SPP1 network signaling across the same FOVs as (A).

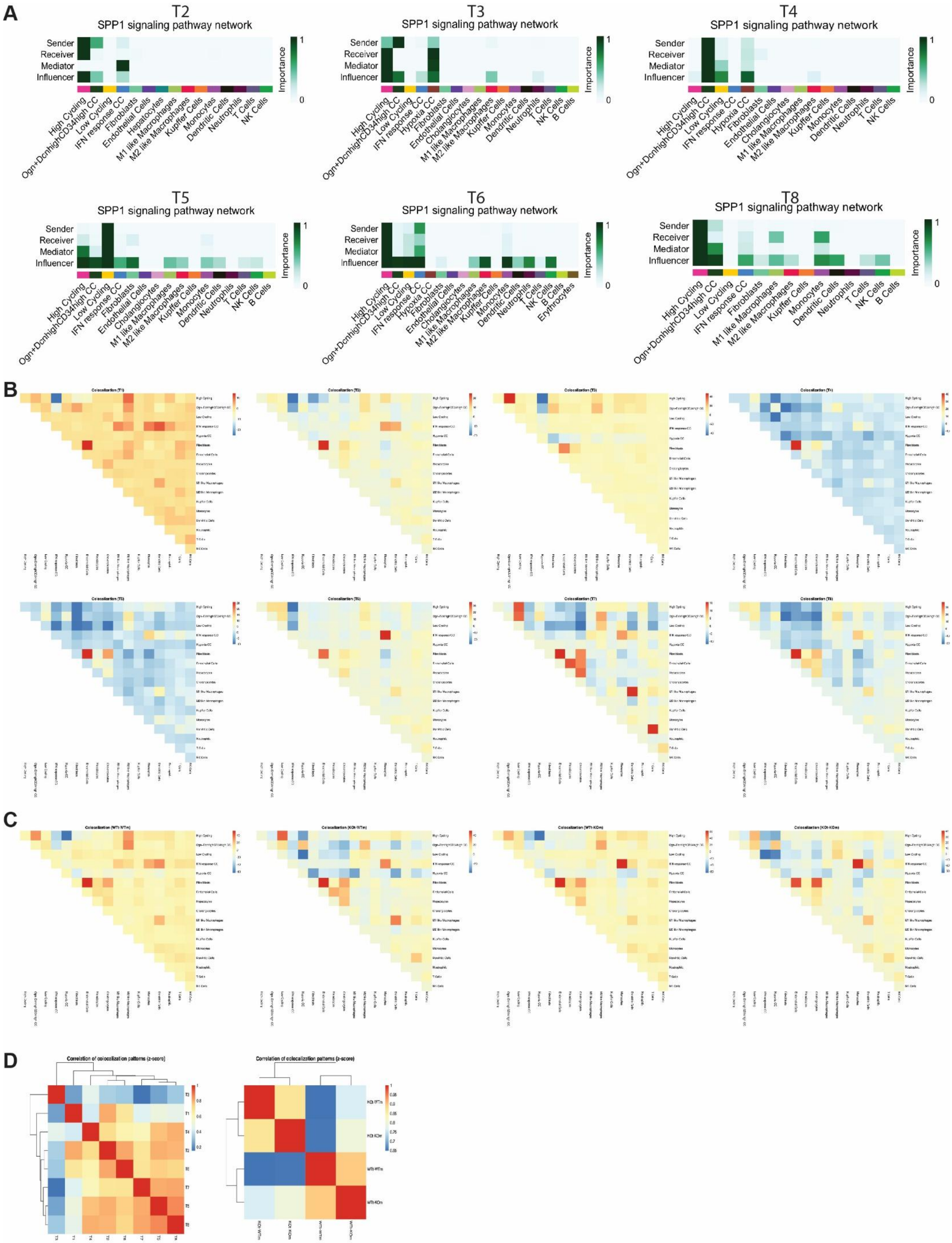
A
T2
SPP1 signaling pathway network
T3
SPP1 signaling pathway network
T4
SPP1 signaling pathway network
T5
SPP1 signaling pathway network
T6
SPP1 signaling pathway network
T8
SPP1 signaling pathway network
Sender
Receiver
Mediator
Influencer
Importance
1
0
High Cycling
Ogn+DcnhighCD34high CC
Low Cycling
IFN response CC
Hypoxia CC
Fibroblasts
Endothelial Cells
Hepatocytes
Cholangiocytes
M1 like Macrophages
M2 like Macrophages
Kupffer Cells
Monocytes
Dendritic Cells
Neutrophils
T Cells
NK Cells
B Cells
Erythrocytes
B
C
D
Correlation of colocalization patterns (z-score)

**Fig S7. SPP1 signaling pathway networks and cell cluster co-localization**

(A) Heatmaps of the SPP1 Signaling network showing the relative importance of each cell group, ranked by the four computed network centrality measures, across the T1-T8 niches.
(B) Correlation heatmaps showing co-localization of cells across tumor niches T1-T8.
(C) Correlation heatmaps showing co-localization of cells across tumor samples (WTt-WTm, KOt-WTm, WTt-KOm, KOt-KOm)
(D) Z-score of correlation of colocalization patterns split by niche (left) and by tumor sample (right)

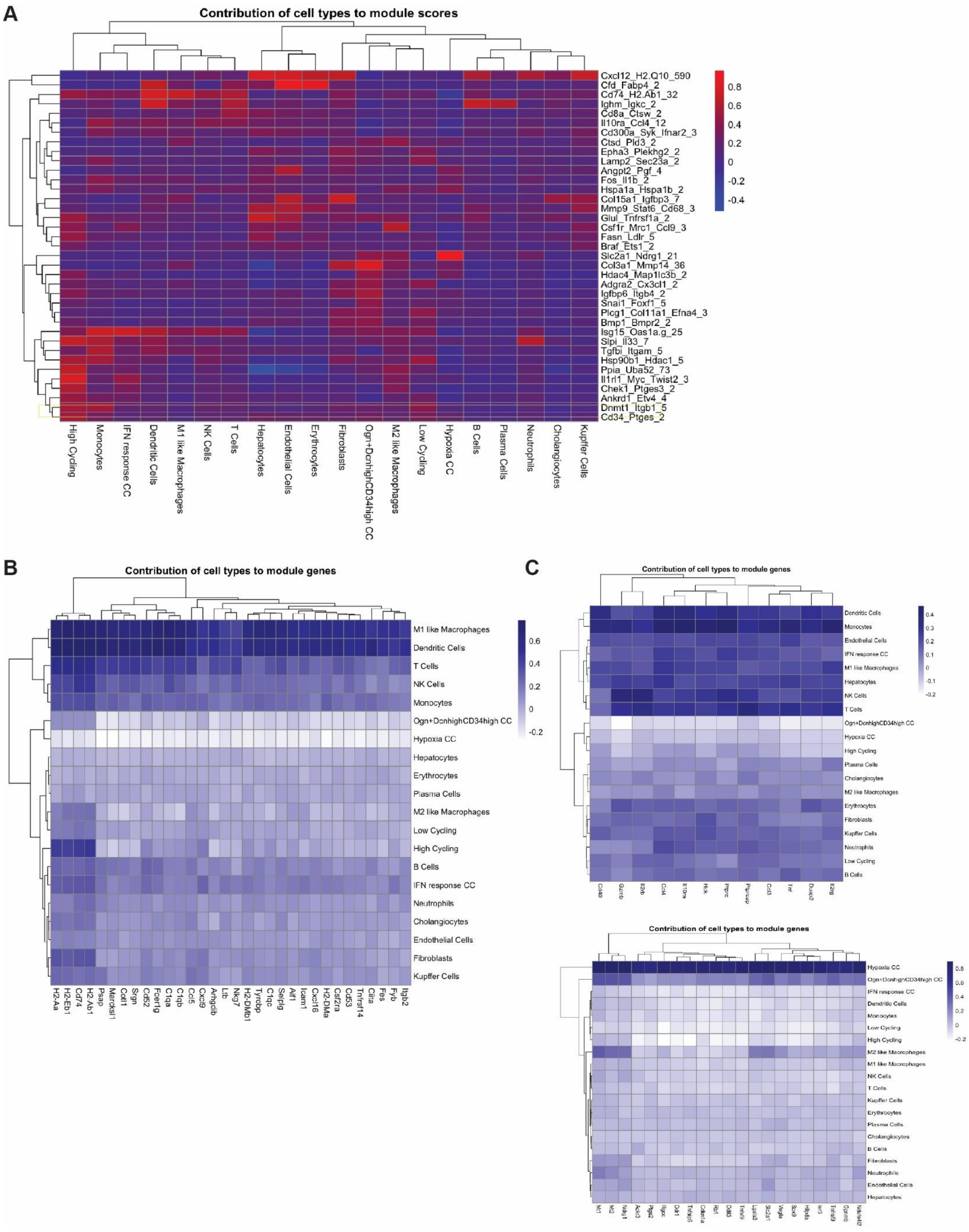

A
Contribution of cell types to module scores
Cxcl12_H2.Q10_590
Cfd_Fabp4_2
Cd74_H2.Ab1_32
Ighm_Igkc_2
Cd8a_Ctsw_2
Il10ra_Ccl4_12
Cd300a_Syk_Ifnar2_3
Ctsd_Pld3_2
Epha3_Plekhg2_2
Lamp2_Sec23a_2
Angpt2_Pgf_4
Fos_Il1b_2
Hspa1a_Hspa1b_2
Col15a1_Igfbp3_7
Mmp9_Stat6_Cd68_3
Glul_Tnfrsf1a_2
Csf1r_Mrc1_Ccl9_3
Fasn_Ldlr_5
Braf_Ets1_2
Slc2a1_Ndrg1_21
Col3a1_Mmp14_36
Hdac4_Map1lc3b_2
Adgra2_Cx3cl1_2
Igfbp6_Itgb4_2
Snai1_Foxf1_5
Plcg1_Col11a1_Efna4_3
Bmp1_Bmpr2_2
Isg15_Oas1a.g_25
Slpi_Il33_7
Tgfbi_Itgam_5
Hsp90b1_Hdac1_5
Ppia_Uba52_73
Il1rl1_Myc_Twist2_3
Chek1_Ptges3_2
Ankrd1_Etv4_4
Dnmt1_Itgb1_5
Cd34_Ptges_2
0.8
0.6
0.4
0.2
0
-0.2
-0.4
High Cycling
Monocytes
IFN response CC
Dendritic Cells
M1 like Macrophages
NK Cells
T Cells
Hepatocytes
Endothelial Cells
Erythrocytes
Fibroblasts
Ogn+DcnhighCD34high CC
M2 like Macrophages
Low Cycling
Hypoxia CC
B Cells
Plasma Cells
Neutrophils
Cholangiocytes
Kupffer Cells
B
Contribution of cell types to module genes
M1 like Macrophages
Dendritic Cells
T Cells
NK Cells
Monocytes
Ogn+DcnhighCD34high CC
Hypoxia CC
Hepatocytes
Erythrocytes
Plasma Cells
M2 like Macrophages
Low Cycling
High Cycling
B Cells
IFN response CC
Neutrophils
Cholangiocytes
Endothelial Cells
Fibroblasts
Kupffer Cells
0.6
0.4
0.2
0
-0.2
H2-Aa
H2-Eb1
Cd74
H2-Ab1
Psap
Marcksl1
Cotl1
Srgn
Cd52
Fcer1g
C1qa
C1qb
Ccl5
Cxcl9
Arhgdib
Ltb
Nkg7
H2-DMb1
Tyrobp
C1qc
Selplg
Aif1
Icam1
Cxcl16
H2-DMa
Csf2ra
Cd53
Tnfrsf14
Ciita
Fes
Fyb
Itgb2
C
Contribution of cell types to module genes
Dendritic Cells
Monocytes
Endothelial Cells
IFN response CC
M1 like Macrophages
Hepatocytes
NK Cells
T Cells
Ogn+DcnhighCD34high CC
Hypoxia CC
High Cycling
Plasma Cells
Cholangiocytes
M2 like Macrophages
Erythrocytes
Fibroblasts
Kupffer Cells
Neutrophils
Low Cycling
B Cells
0.4
0.3
0.2
0.1
0
-0.1
-0.2
Cd40
Gzmb
Il2rb
Ccl4
Il10ra
Hck
Ptprc
Ptprcap
Ccl3
Tnf
Dusp2
Il2rg
Contribution of cell types to module genes
Hypoxia CC
Ogn+DcnhighCD34high CC
IFN response CC
Dendritic Cells
Monocytes
Low Cycling
High Cycling
M2 like Macrophages
M1 like Macrophages
NK Cells
T Cells
Kupffer Cells
Erythrocytes
Plasma Cells
Cholangiocytes
B Cells
Fibroblasts
Neutrophils
Endothelial Cells
Hepatocytes
0.8
0.6
0.4
0.2
0
-0.2
Mt1
Mt2
Ndrg1
Ackr3
Ptgs2
Rgcc
Ddr1
Tnfaip6
Cdkn1a
Rb1
Ddit3
Tnfsf9
Lgals3
Slc2a1
Vegfa
Sox9
Hilpda
Ier3
Tnfrsf9
Gpnmb
Ndufa4l2

**Figure S8. Contributions of cell types and module score.**

(A) Involvement of each cell type in the top-ranked spatially correlated gene modules. The column indicates cell type, and the row represents gene modules identified by InSituCor package. The number at the end of module name indicate the number of genes in each module.

(B-C) Contribution of each cell type to the genes in representative spatially correlated modules such as CD74_H2-Ab1_32 (B), Il10ra_Ccl4_12 (C), and Slc2a1_Ndrg1_21 (D)
